# Genomic and transcriptomic analyses reveal a tandem amplification unit of 11 genes and mutations of mismatch repair genes in methotrexate-resistant HT-29 cells

**DOI:** 10.1101/2020.02.26.965814

**Authors:** Ahreum Kim, Jong-Yeon Shin, Jeong-Sun Seo

## Abstract

*DHFR* gene amplification is present in methotrexate (MTX)-resistant colon cancer cells and acute lymphoblastic leukemia. However, little is known about *DHFR* gene amplification due to difficulties in quantifying amplification size and recognizing the repetitive rearrangements involved in the process. In this study, we have proposed an integrative framework to characterize the amplified region by using a combination of single-molecule real time sequencing, next-generation optical mapping, and chromosome conformation capture (Hi-C). Amplification of the *DHFR* gene was optimized to generate homogenously amplified patterns. The amplification units of 11 genes, from the *DHFR* gene to the *ATP6AP1L* gene position on chromosome 5 (~2.2Mbp), and a twenty-fold tandemly amplified region were verified using long-range genome and RNA sequencing data. In doing so, a novel inversion at the start and end positions of the amplified region as well as frameshift insertions in most of the *MSH* and *MLH* genes were detected. These might stimulate chromosomal breakage and cause the dysregulation of mismatch repair pathways. Using Hi-C technology, high adjusted interaction frequencies were detected on the amplified unit and unsuspected position on 5q, which could have a complex network of spatial contacts to harbor gene amplification. Characterizing the tandem gene-amplified unit and genomic variants as well as chromosomal interactions on intra-chromosome 5 can be critical in identifying the mechanisms behind genomic rearrangements. These findings may give new insight into the mechanisms underlying the amplification process and evolution of drug resistance.

## Introduction

Gene amplification, the triggering of an abnormal copy number increase in a specific region of the genome of an organism growing in a selective condition, is associated with overexpression of oncogenes such as *MYC, MYCN*, and *ERBB*, which engender abnormal cell proliferation and replication^1, 2, 3^. These genes undergo amplification much more frequently than would be caused by a genomic mutation event in mammalian cells, since the rate of gene amplification is greater than the mutation rate ^4^; thus, such amplication has great tumorigenic potential^5^.

However, no molecularly targeted agents have been specifically developed to prevent gene amplification because of the associated chromosomal complexity and technical limitations. It is therefore critical to find predictive and prognostic biomarkers for gene amplification to develop specific medicines that will improve patient outcomes and optimize therapeutic decisions^6, 7^.

In addition, gene amplification is an indicator of a drug-resistant sample in cancer and healthy cells^8^, so it will be important to identify the genetic features or pathways that promote amplification in tumors. These mechanisms might serve as therapeutic targets that can prevent drug resistance and arrest or eradicate a tumor^9^. However, the molecular mechanisms that contribute to high gene copy numbers are completely unknown, since sequence alignment and assembly programs that rely on short reads are not equipped to deal with genomic rearrangements and repetitive sequences^10, 11^.

*DHFR* gene amplification at chromosome 5 has been a hallmark of methotrexate (MTX) responsiveness and resistance in colon cancer cells and acute lymphoblastic leukemia. MTX is an antifolate drug that inhibits dihydrofolate reductase (*DHFR*) by preventing DNA synthesis and cell division^12, 13, 14^.

The amplified *DHFR* gene generates two major DNA segments consisting of extrachromosomal double-minutes (DMs) and intrachromosomal homogenously staining chromosome regions (HSRs); however, the molecular mechanisms behind these amplified region products and suitable methods for their detection and characterization are still unkown. Currently, these segments are detected only in advanced-stage tumors and accompany more repetitive and complicated sequences to assemble^15, 16 17^.

In this study, single-molecule real time (PacBio SMRT) sequencing, optical genome mapping (BioNano Genomics; read size: ~10Kb), and high throughput chromosome conformation capture (Hi-C) for inter- and intra-chromosomal interactions are used to identify relevant repetitive rearrangements with amplified segments and interpret gene amplification mechanisms in a MTX-resistant colon cancer cell line (HT-29) (Fig. 1), which has a heterogeneously amplified genome.

**Figure 1.**
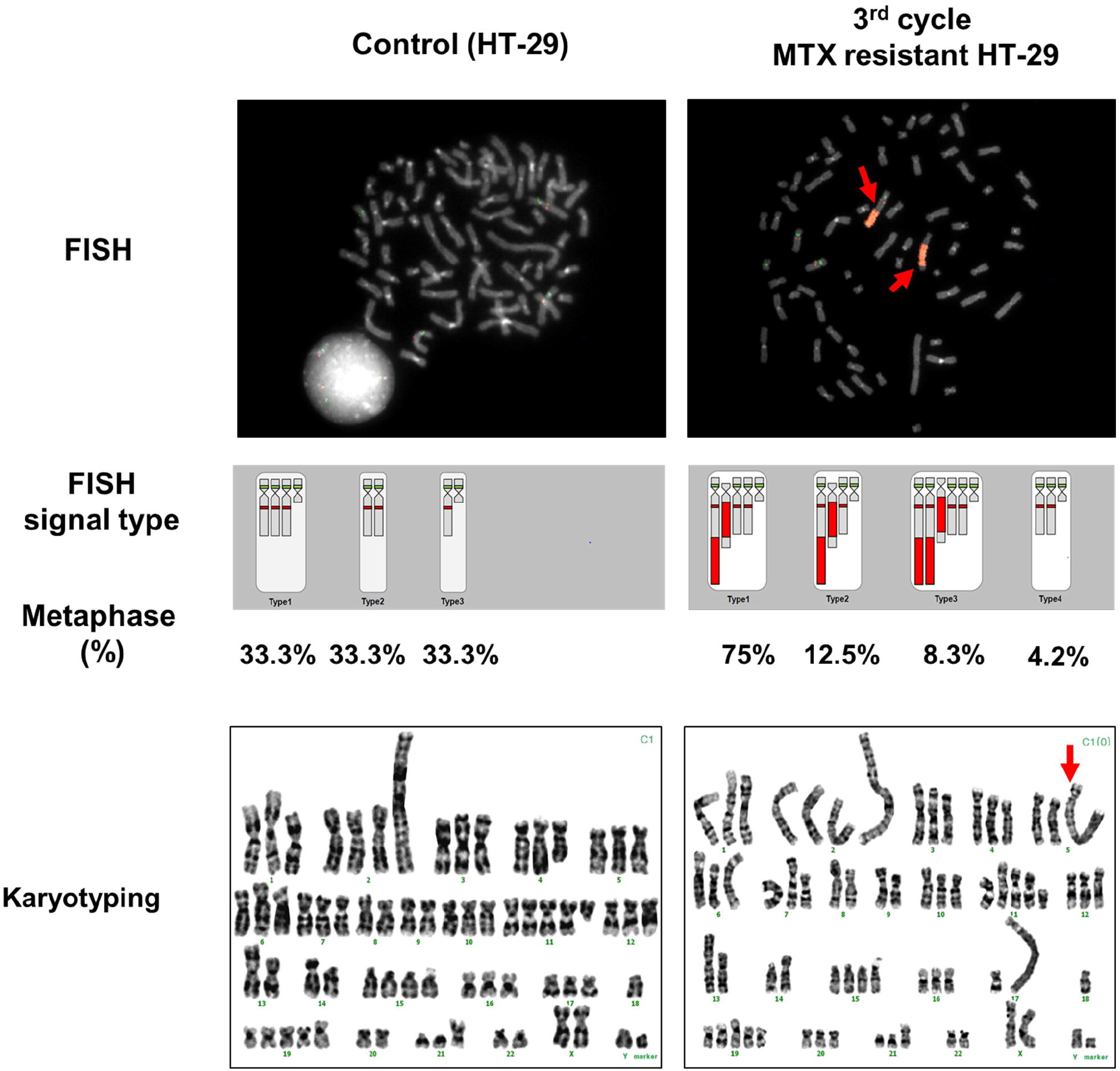
Visualization of *DHFR*-amplification patterns using FISH and karyoptyping between MTX-resistant clones (C1-2-4) and controls. A subculture (C1-2-4) of the C1-2 clone was visualized via fluorescent in situ hybridization (FISH), at 1000x magnification, and the FISH signal type and percentage at metaphase were compared between controls and MTX-resistant clones (C-1-2-4). All chromosomes were karyotyped and abnormal chromosomal shapes were detected at the 5q and 17q arms, compared to controls, as indicated by the red arrow.

These techniques allowed for accurate quantification of amplification size and identification of drastic differences in chromosomal abnormalities and structural variants compared to MTX-sensitive sample, which are difficult to extensively analyze using short-read sequencing and fluorescence in situ hybridization (FISH). Accordingly, this study contributes to an understanding of basic genomic principles, impacts of genetic rearrangements on cancer cells, and by extension, drug resistance mechanisms.

## Results

### Determination of MTX-resistant HT-29 cells

While generating MTX-resistant HT-29 cells (selected clone: C1-2) from singe-cell selection and MTX sensitization as previously described^15^, dramatic morphological changes in the cells themselves were detected in the form of rounded and circular shapes at the 1^st^ cycle of sensitization as well as rod and irregular shapes at the 2^nd^ cycle (**Supplementary Fig. 1**). The original shapes were again observed at the 3^rd^ cycle of sensitization, which might indicate that the HT-29 cells were becoming resistant to MTX and rapidly grew up under high MTX concentration.

As expected, *DHFR* expression, which was normalized by the *B2M* housekeeping gene, was steadily increased from the 1^st^ to 3^rd^ cycle, as the HT-29 clones became resistant to MTX (**Supplementary Fig. 2a**). After confirming these morphological changes and increased *DHFR* expression, the *DHFR* copy number was measured in several clones.

The clones’ cycle quantification values (Ct) were used to compute the rate of copy number. These values dropped from 26.04 to 19.99, as the number of cycles increased (**Supplementary Table 1**). Interestingly, there was a dramatic increase in the *DHFR* copy number between the 1^st^ (0.97 copies) and 2^nd^ (54.83 copies) cycles (**Supplementary Fig. 2b**).

**Table 1.** The comparison of detected structural variants between control and MTX resistant HT-29.

| No. of Events<br>(median size) | All chromosomes |  | Chromosome 5 |  |
| --- | --- | --- | --- | --- |
|  | Control (HT-29) | MTX resistant HT-29 | Control (HT-29) | MTX resistant HT-29 |
| Deletion | 13(4472) | 14(2917) | 2(4425.5) | 3(1514) |
| Duplication | 1(1004) | 5(13534) | 0 | 1(371675) |
| Inversion | 0 | 4(331155.5) | 0 | 3(328697) |
| Inverted Duplication | 8(47) | 15(77) | 0 | 3(1529) |
| Translocation | 45 | 50 | 4(chr3 to chr5) and<br>(chr5 to chr12) | 2(chr5 to chr12) |

The structural variants ( deletion, duplication, inversion inverted duplication, and translocation) were categorized by using Sniffles from sorted PacBio output derived from BWA-MEM over all chromosomes and chromosome 5 in control(HT-29) and MTX resistant HT-29 cells. The median size of split reads is shown in parentheses

Based on these results, we ascertained that a specific time period and set of conditions were required for clones to survive in the presence of MTX, and the amplified *DHFR* gene was an indicator of MTX resistance, as previously described^12^. Also, it was confirmed that the increase in *DHFR* gene expression was not proportional to the increase in *DHFR* gene copy number in each cycle^18^.

### Validation of gene amplification in MTX-resistant HT-29

After quantifying the *DHFR* gene copy number, the amplified *DHFR* gene at the 5q arm was visualized via fluorescent in situ hybridization (FISH) for a MTX-resistant clone (C1-2) and control sample. The *DHFR* gene region at the 5q arm was abnormally stretched compared to that of the control, at the 2^nd^ and 3^rd^ cycle, as expected (**Supplementary Fig. 3**).

The amplified *DHFR* gene patterns of C1-2 cells in metaphase revealed that the amplified region had a painting signal, and FISH signal patterns were highly heterogeneous. A total of 9 signal-amplified patterns and two major patterns accounted for 44% and 28% of abnormally DHFR gene amplified patterns, respectively (**Supplementary Fig. 4**). Even if the cells suffered from the same condition, the MTX effects and genetic status were different in each cell. This would significantly hinder technical analysis of the alignment and analysis of amplified patterns^19, 20^. Therefore, we had to optimize homogenously amplified patterns before sequencing by performing a serial clone selection and sub-culturing.

The optimized C1-2 clone (C1-2-4), which was robust in the presence of MTX, was visualized using FISH. Four different types of gene amplification pattern were detected; two patterns had amplified *DHFR* genes at two q arms (75 % and 12.5 %), one pattern had amplified DHFR genes at three q arms (8.3 %), and one completely lacked amplified *DHFR* genes at q arms (4.2 %) (Fig. 1). Overall, ~96% of the cells had a *DHFR*-amplified region.

A majority of these strains was resistant to MTX and had high amplification of *DHFR* genes at a 5q arm. However, each MTX-resistant C1-2-4 cell had a different *DHFR* gene amplification pattern and copy number, as well as heterogeneous genetic status in the presence of MTX, even if the clone was generated from a single MTX-resistant cell^21^.

Additionally, MTX-resistant HT-29 and control samples were karyotyped to accurately detect the amplified region. The C1-2-4 clone had a homogeneously staining region (HSR) at the chromosome 5 q arm, as previously described^22^, in addition to an abnormal stretch of the 17 q arm. This result confirmed that the aneuploidy and chromosomal instability of cancer samples could cause the many characteristic changes in chromosome copy number and karyotype diversity ^23^. It was previously found that chromosome 17 q arm amplification could be detected because of the genetic instability in colorectal cancer^24^.

### Analysis of the structural variants and amplification unit

After confirming the HSR on the amplified *DHFR* region, the five genomic structural variants (deletion, duplication, inverted duplication, translocation, and inversion) and amplified units of MTX-resistant cells were analyzed to identify which genes and structural variants were involved in gene amplification. The log_2_ ratio of segmented coverage over whole chromosomes between the control and MTX-resistant sample was compared. A high segmented coverage in the MTX-resistant sample, compared to that of the control sample, was observed in chromosome 5 (Fig. 2a). The genes for which the segmented coverage was bigger than 20 were identified and annotated by position to identify the exact amplified region that included the *DHFR* gene.

**Figure 2.**
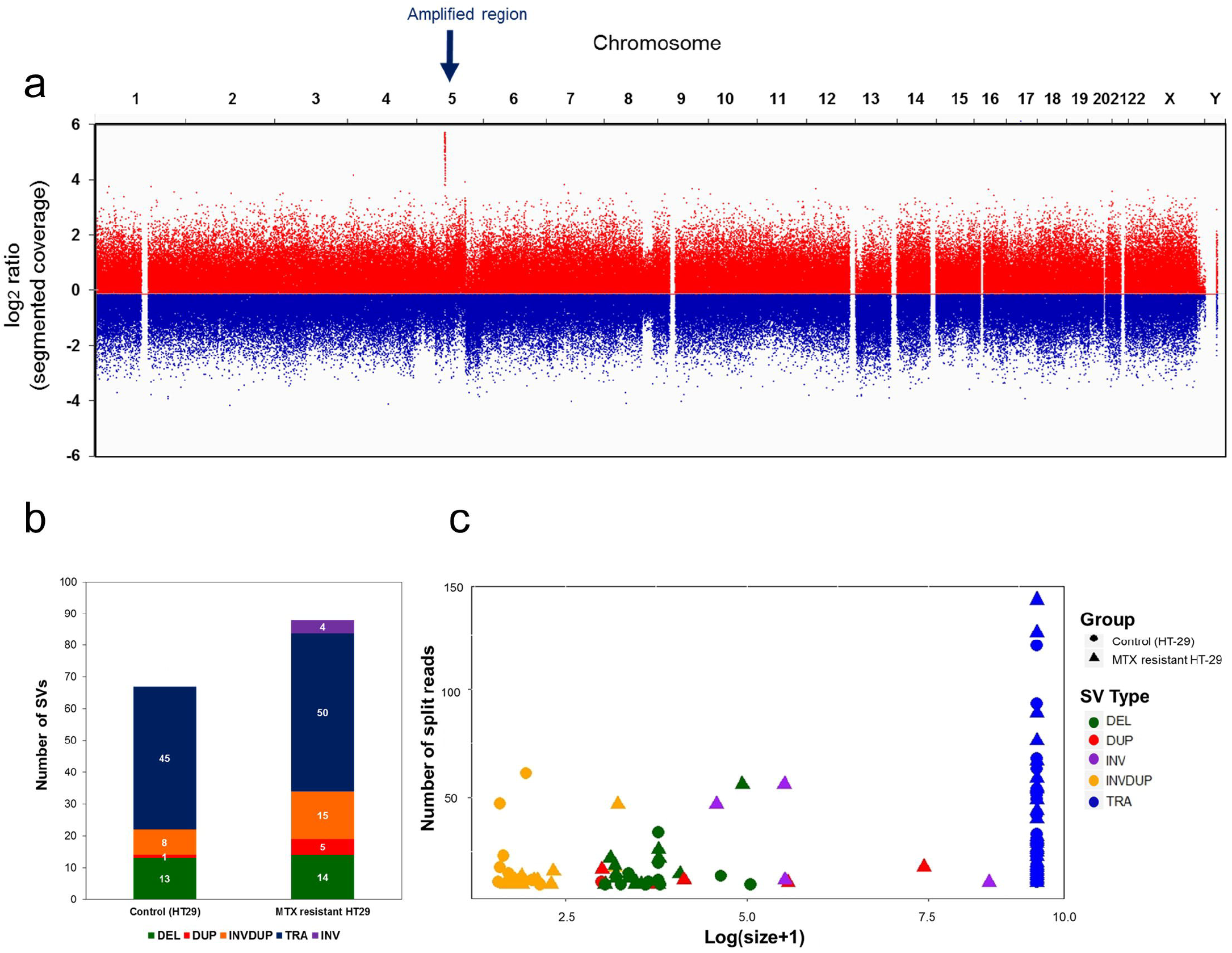
Detection and characterization of SVs in MTX-resistant HT-29 cells. (a) The log2 ratio of segmented coverage across all chromosomes was compared between control and MTX-resistant HT-29 cells. The amplified region on 5q in MTX-resistant HT-29 cells is indicated by the blue arrow. (b) Five genomic variants (deletion, duplication, inverted duplication, translocation, and inversion) in MTX-resistant sample and control were analyzed and visualized with a bar graph, wherein the number of variants was counted and compared between the two samples. (c) In the scatter plot, the size of variants (log(size+1)) and depth of split-reads were plotted and compared between control and MTX-resistant HT 29 cells. The group and variant type are delineated by a different shape and color.

The total number of genomic variants in the MTX-resistant sample was bigger than in the control, and the size of variants such as duplicates and inversions was bigger in the MTX-resistant sample, which had a large number of split reads (Fig. 2b and 2c). Additionally, more variants for each structural variant were detected in MTX-resistant HT-29 cells, compared to those of the control.

The novel structural variants on chromosome 5 were selectively compared between control and MTX-resistant HT-29 cells. One duplication (median size of split reads: 371,675), three inversions (median size of split reads: 328,697), and three inverted duplications (median size of split reads: 1,529) were detected, while there were no such variants detected in the control sample (Table 1). Moreover, the number of split reads for HT-29 cells was bigger than the average coverage (10X), and most copy number variation (CNV) categories matched, which indicated that the detected structural variants had been accurately detected (**Supplementary Table. 2**). Finally, although the number of translocations on chromosome 5 was decreased in MTX-resistant sample, there were more detected inter-chromosomal genomic rearrangements in MTX-resistant HT-29 sample (**Supplementary Fig. 5**).

The position of the amplified region on the chromosome 5 q arm was between 80M to 83M, which included the *DHFR* gene as the start-point and the *ATP6AP1L* gene. The segmented coverage was approximately 197X, between 80.6M and 82.8M (~2.2M) (Fig. 3a). The MTX-resistant sample had an approximately twentyfold longer amplified region than that of the control. This amplified size was inferred from the designed coverage of long read sequencing, which was 10X that of the control sample, but the MTX-resistant sample had abnormally high coverage (~197X) for only this region.

**Figure 3.**
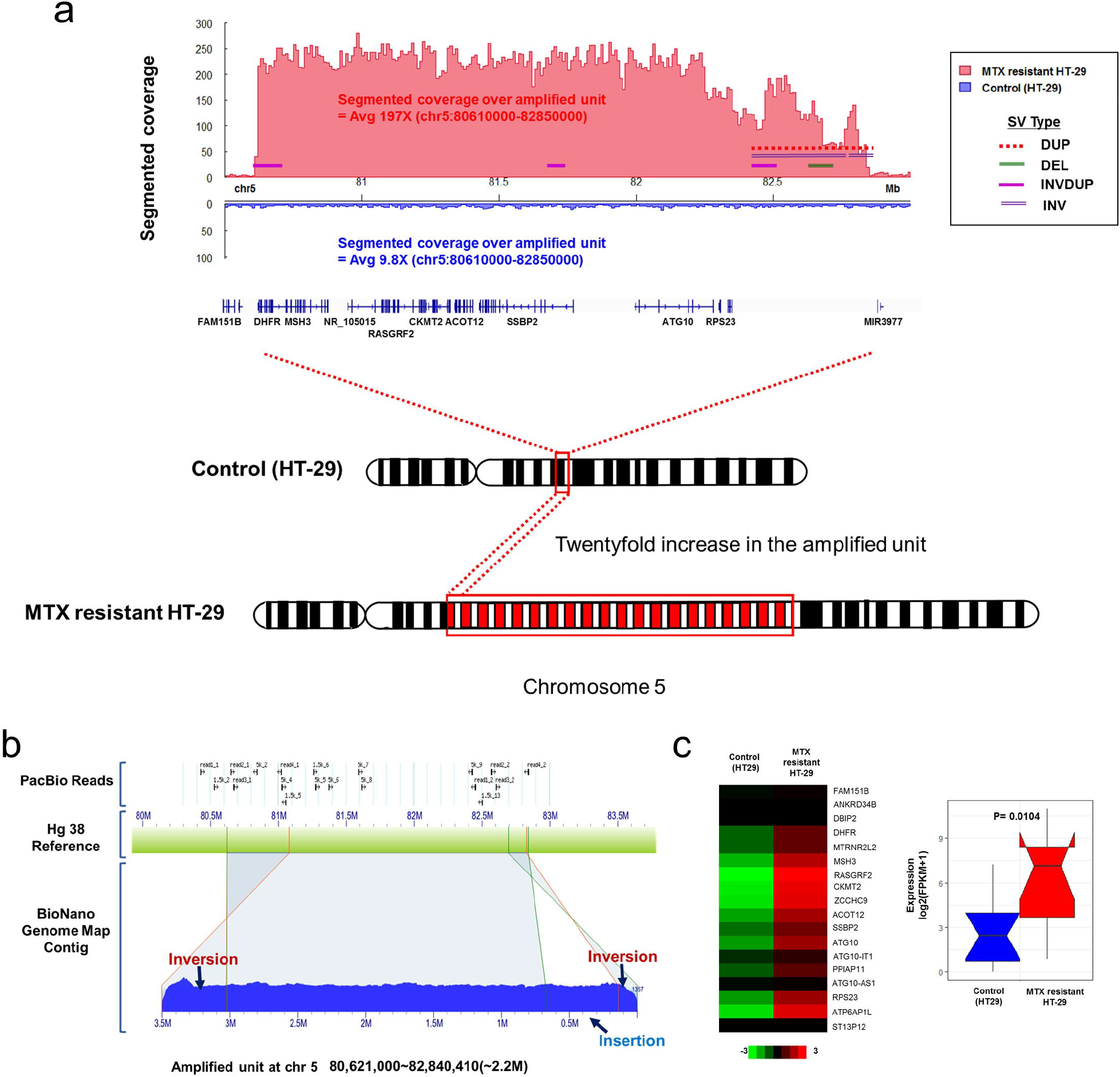
Detection of amplified units and structural variants over the amplified region in MTX-resistant HT-29 cells. (a) The segmented coverage and genomic variants were compared and visualized between control and MTX-resistant HT-29 cells across the amplified region on chromosome 5 (80,610,000-82,850,000). (b) The BioNano contig and PacBio long-reads matched up with the reference (hg38); several genomic rearrangements are indicated by blue arrows. (c) The heatmap depicts the gene expression (FPKM) in amplified units between MTX-resistant HT-29 cells and control. High expression in a specific region was indicated by a red arrow. The statistical significance (p-value) was computed using the Mann Whitney test.

Interestingly, there were both forward and reverse strands in the MTX-resistant sample that were not found in the control sample. These had only forward strands and small deletions in our defined amplified unit (**Supplementary Fig. 6**). Previous studies have shown that forward DNA synthesis is preferred during replication^25^; however, this result indicated that the replication was performed in the both forward and reverse direction in the MTX-resistant samples. A study by Kimura et al. (2016) found that DNA can be replicated in the reverse direction by a backward enzyme such as Thg1-like proteins (TLPs) to efficiently synthesize both chains^26^.

Overall, the amplification regions included the 11 genes, from *DHFR* to *ATP6AP1L*, which were tandem gene amplifications of this region, which had inversions or inverted duplications at the end of the amplification unit. The tandem repeats of several genes on chr5 (2.2Mbp) were initiated and terminated by inversion on the specific sequence. Such an inversion (chr5:80,618,750-80,631,409) at the start point included the *LINC01137, DHFR, CTC325J23.2*, and MTRNR2L2 genes (**Supplementary Fig. 7**).

### Optical genome mapping over the amplified region

The region of tandem amplification (80.6Mbp – 82.8Mbp) on chromosome 5 was further analyzed using BioNano genome optical mapping, due to a lack of coverage and mapping over the whole range of the amplified region using other methods. The contigs of genome mapping mapped in complex ways to whole chromosomes, except for the amplified region, with high coverage of approximately 200x (**Supplementary Fig. 8**). Additionally, the gene-amplified unit had inversions at both the start and endpoints of the amplified region, as expected, but there was a newly identified insertion at the endpoint of the amplified region (Fig. 3b).

These inversions were certainly associated with the amplification mechanism and seemed to assist with and even initiate tandem repeat amplification, as previously reported^27^. The identified inverted repeat could stimulate the formation of a large DNA palindrome after breakage of an adjacent DNA double-strand^28^. This suggests that short inversion at the start and endpoints in MTX-resistant HT-29 cells could play an important role in the initiation of gene amplification.

### Expression levels of MTX-resistant sample

There was extremely high gene expression within the identified amplification region, which matched with the high coverage of the amplified unit in our long-read sequencing data (**Supplementary Fig. 9**). The read coverage from long-read sequencing of the amplified region in the MTX-resistant sample was approximately ten times higher than that of the control. Similarly, the log_2_(FPKM+1) expression level from the *DHFR* to *ATP6AP1L* gene in MTX-resistant HT-29 cells was 5x that of control for *DHFR* and 122x that of control for *RASGRF2* (*P* = 0.0104 via the Mann-Whitney test; Fig. 3c).

### Identifying novel mutations and their impacts on gene amplification

In order to find relevant mutations on the amplification mechanism, single-nucleotide variants (SNVs) in both samples were identified using transcriptome sequencing data. There were more total exonic mutations in MTX-resistant HT-29 cells (13,982) compared to those of control sample (13,310) across all chromosomes, and 18 more exonic mutations were detected in MTX-resistant HT-29 cells on chromosome 5 only (**Supplementary Tables 3 and 4**).

After filtering the synonymous SNVs, there were a few additional non-synonymous mutations (non-synonymous SNVs, frameshift deletions, frameshift insertions, stop-gains, and stop-losses) in MTX-resistant HT-29 cells than those for control, across all chromosomes (**Supplementary Fig. 10**). Some non-synonymous mutations came from chromosome 5, in which the number and percentage of frameshift insertions noticeably increased in MTX-resistant sample (18.1%) than in control (4.3 %) (**Supplementary Fig. 4a**). The most frequently inserted nucleotides in frameshift insertions were thymine (73 %) and adenine (23 %).

This might explain why the detected frameshift thymine and adenine insertions in mRNA on chromosome 5 concurrently occurred with *DHFR* gene amplification, which conferred an ability to survive MTX exposure. Moreover, novel frameshift insertions -- either adenine (A) or thymine (T) -- were located on *MSH3* and *MSH6* genes as well as *PMS1* and *PMS2* genes in MTX-resistant HT-29 sample only (**Supplementary Table. 5**). Expression of these genes, except for *MSH3*, decreased in the mutated and MTX-resistant HT-29 cells, compared to control sample (Fig. 4b).

**Figure 4.**
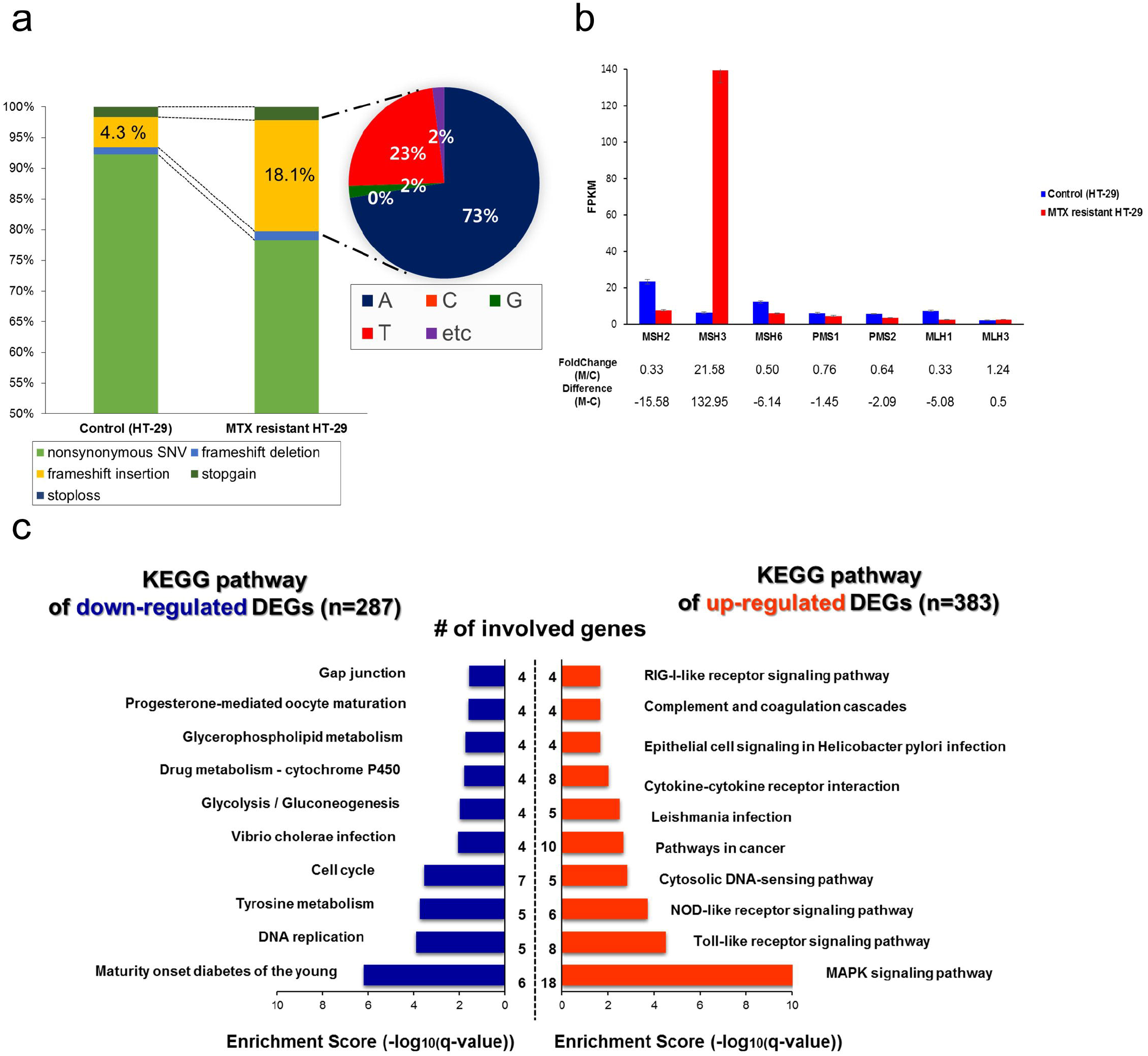
Comparison of non-synonymous mutations and MMR expression levels. (a) Non-synonymous mutations in chromosome 5 (non-synonymous SNV, frameshift deletion, frameshift insertion, stop-gain, and stop-loss) were compared between MTX-resistant HT-29 cells and control. Each portion of the four nucleobases in the frameshift insertions was indicated in the MTX-resistant HT-29 cells. (b) Gene expression (FPKM) of mutS homologs and mutL homologs was computed and compared. The change and difference between MTX-resistant HT-29 cells (M) and control (C) are indicated below the bar graph. (c) The top 10 enriched KEGG gene sets for up-regulated and down-regulated differentially expressed genes were visualized via an enrichment score (-log(qvalue)).

The *MSH3* and *MSH6* genes belong to mismatch repair (MMR) ^29^ gene families and are known to play an important role in repairing DNA for cell division as well as cooperatively suppressing intestinal tumors^30^. MLH genes (*PMS1*) as well as the MSH genes (*MSH1 - 6*) are significantly correlated with colon cancer; mutations in these genes could cause predisposition and susceptibility to Lynch syndrome, in conjunction with colon cancer^31, 32, 33^.

Therefore, novel frameshift insertions in these genes could prevent mismatch repair functionality and tumor suppression in the presence of MTX, and stimulate the rapid progression of gene amplification and MTX-resistance. The molecular explanation for this tandem-gene amplification mechanism could be a malfunction of MMR pathways^34^. It was previously known that *MSH3* is concurrently amplified with the *DHFR* gene due to their proximity to each other in MTX-resistant cells. An imbalance in expression of mutS homologs could result in the malfunctioning of base-base mismatch repair and cause genetic instability as well as confer resistance to the cytotoxic effects of MTX^35^. Also, chromothripsis events by massive clustered genomic rearrangements might contribute to tandem gene amplification and inactivation of MMR genes under MTX condition^36^.

### Identifying differentially expressed genes (DEGs)

To determine which genes and gene sets are involved in MTX-resistance, we analyzed differentially expressed genes and enriched gene sets (Fig. 4c). A total of 383 up-regulated and 287 down-regulated DEGs were identified in MTX-resistant HT-29 cells and compared to those of control sample (**Supplementary Table. 6**).

Through enrichment analysis of KEGG gene sets, the up-regulated DEGs included *IL1B, MAPK11, JUN, MAP3K8, IL8,and CASP1* in MTX-resistant HT-29 cells. The affected signaling pathways included MAPK, toll-like receptor, and NOD-like receptor signaling. Interestingly, the down-regulated DEGs such as *MAD2L1, CCNA2, MCM2, MCM4, FEN1*, and *CDK6* were enriched in DNA replication, tyrosine metabolism, and cell cycle pathways, which are commonly up-regulated in colon cancer^37^.

As previously reported, the down-regulated DEGs of MTX-resistant osteosarcoma cell lines were enriched in the mitotic cell cycle, cell cycle, and DNA replication pathways, which were also down-regulated in our MTX-resistant colon cancer cells. This result might explain the role of MTX in inhibiting dihyrofolate reductase (*DHFR*) and preventing tumor cells from proliferating in both cases^38, 39^. In addition, down-regulated expression of MMR genes, which affects G2/M cell cycle arrest and apoptosis, might prevent proper checkpoint and cell-death signaling; therefore, this phenomenon could contribute to the malfunctioning of DNA replication^40^.

Still, this does not explain which mechanism is associated with *DHFR* gene amplification in MTX-resistant colon cancer cells and cannot be used to identify MTX targets or other intracellular pathways and folate metabolism markers^41^.

### Chromosomal interactions and their topologically associated domains

Genome-wide Intra-chromosomal interactions were identified and compared between MTX-resistant HT-29 cells and control, at 5kb resolution (**Supplementary Fig. 11**). A high interaction frequency was apparent due to a clear long red line on the amplified region only (5q14.1 to 5q14.2) in the MTX-resistant HT-29 cells. This interaction pattern was similar to those observed when amplification occurred in tumor samples in a previous study^42^(**Supplementary Fig. 12**).

The topologically associating domains (TADs) and several chromosomal rearrangements at chromosome 5 were identified to visualize conformations and interactions on intra or inter-chromosomes within the amplified region, as well as to detect unforeseen chromosomal rearrangements at 500kb resolution (Fig. 5a). Frequent intra-chromosomal interactions were observed in the amplified region (chr 5:80.6M-82.8M), and there were several newly identified TADs in the middle and endpoint of this region compared to those of control with high adjusted interaction frequencies (adjusted M) and an adjusted p-value < 0.05 (**Supplementary Table 7**).

**Figure 5.**
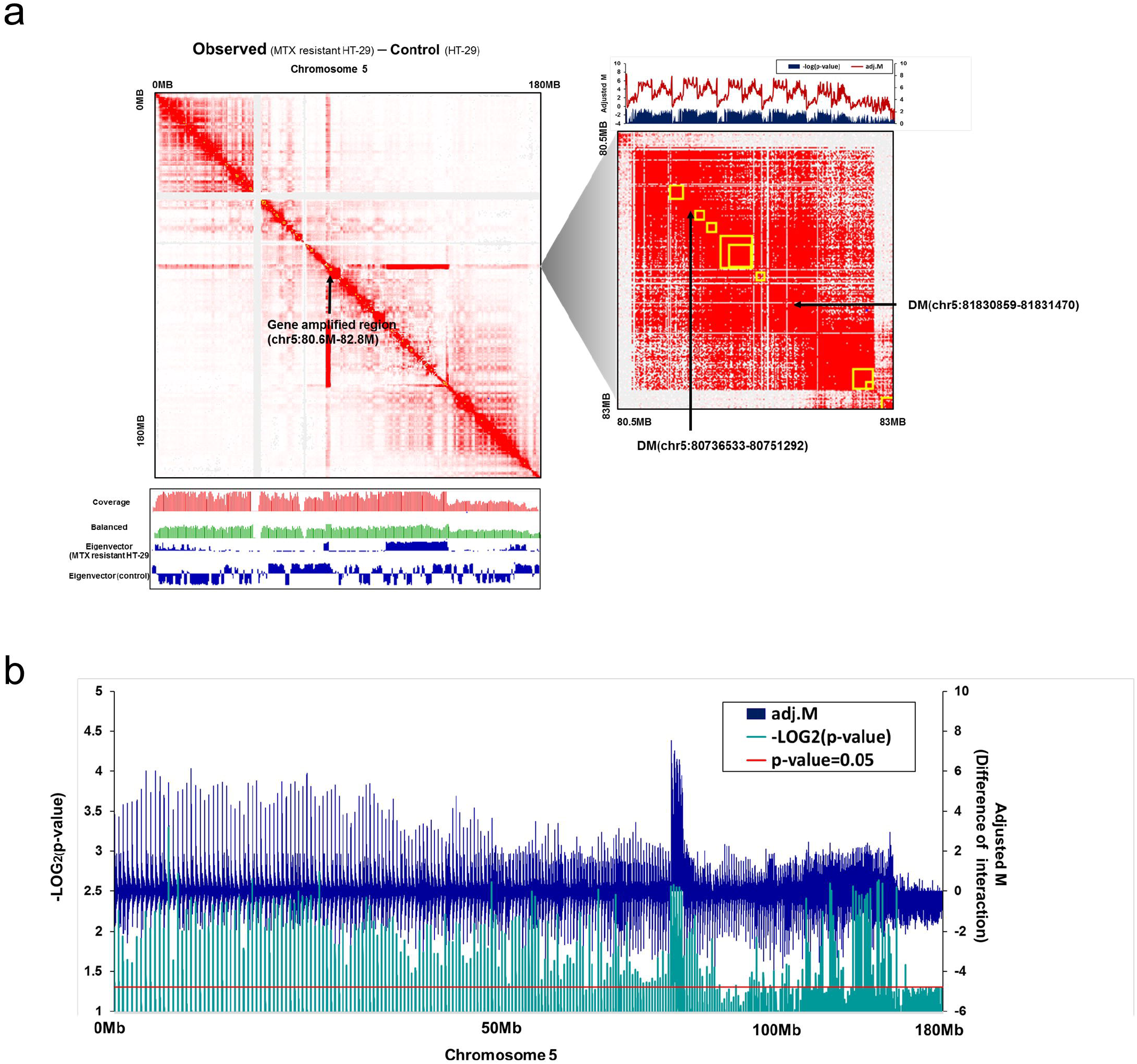
Topologically associating domains (TADs) on chromosome 5 and adjusted interaction frequencies. (a) Intra-chromosomal interactions at chromosome 5 (MTX resistant HT-29 – control) are visualized with the coverage and eigenvectors (left), and the newly identified topologically associating domains (TADs) on the duplicated region are indicated by a yellow box, at 500kb resolution (right). (b) The difference in adjusted interaction frequencies (adjusted M) between MTX-resistant and control samples is plotted on a –log2 scale (p-value). Statistical significance (p-value=0.05) is indicated by the red line.

Interestingly, interactions were also frequent within this region between 109Mbp to 138Mbp, and more TADs and high eigenvector values were observed. These delineated compartmentalization in Hi-C^43^ and indicated that both the amplified region and an adjacent region had both frequent intra-chromosomal interactions and frequent contacts, and it seemed that the entire 5q region upregulated the amplification mechanism.

In order to compare the computed intra-chromosomal interactions between MTX-resistant HT-29 cells and control samples, the difference in adjusted interaction frequencies (adjusted M), with a p-value at 500Kb resolution, was analyzed, and the differentially interacting genomic regions at chromosome 5 were identified in the amplified region (p-value < 0.05; Fig. 5b). The adjusted M values were lower in the region from 109Mbp to 138Mbp than those in the amplified region, but this region still had a significant p-value and higher interactions compared to those in other positions. Therefore, the chromosomal structure, from the start position of the amplified unit (80Mb) to its endpoint on 5q, could have a complex network of spatial contacts to harbor gene amplification.

The relative copy number in each chromosome was estimated from Hi-C data, using chromosome 2 as a reference. Specifically, the relative copy number in chromosome 5 was significantly higher in MTX-resistant HT-29 cells compared to control, over whole chromosomes except for chromosome 17, which also had a stretched structure like chromosome 5 (**Supplementary Fig. 13**).

In addition, chromosomal rearrangements such as those that would result in DMs only were extrachromosomal DNA stretches that harbored the amplification of oncogenes that were involved in drug resistance. These rearrangements were detected in the amplified region via Hi-C data, and were not observed in the FISH data. Hi-C data was utilized because it could reveal unforeseen chromosomal rearrangements and copy numbers in highly amplified regions. However, this result should be confirmed using other techniques, since it was not clear how to distinguish between DMs and HSRs due to their similar structure.

Overall, tandem gene amplifications of several genes on chr5 (2.2Mbp) in long read sequencing data were confirmed via gene expression and gene mapping as well as HI-C data. More complicated interactions on intra-chromosome 5 can be critical factors in identifying the hotspots of spatial contacts across an amplified region.

### The mechanisms behind tandem gene amplification

We have developed experimental and computational workflows to detect and analyze the mechanisms behind gene amplification in MTX-resistant HT-29 cells through breakage-fusion-bridge (BFB) cycles, as previously reported^44^ (Fig. 6 and **Supplementary Fig. 14**). Before gene amplification occurred, frameshift insertions in *MSH* and *MLH* genes across several chromosomes were caused by MTX toxicity, which resulted in a genetic predisposition and dysregulation of mismatch repair pathways in the presence of MTX^16, 45^.

**Figure 6.**
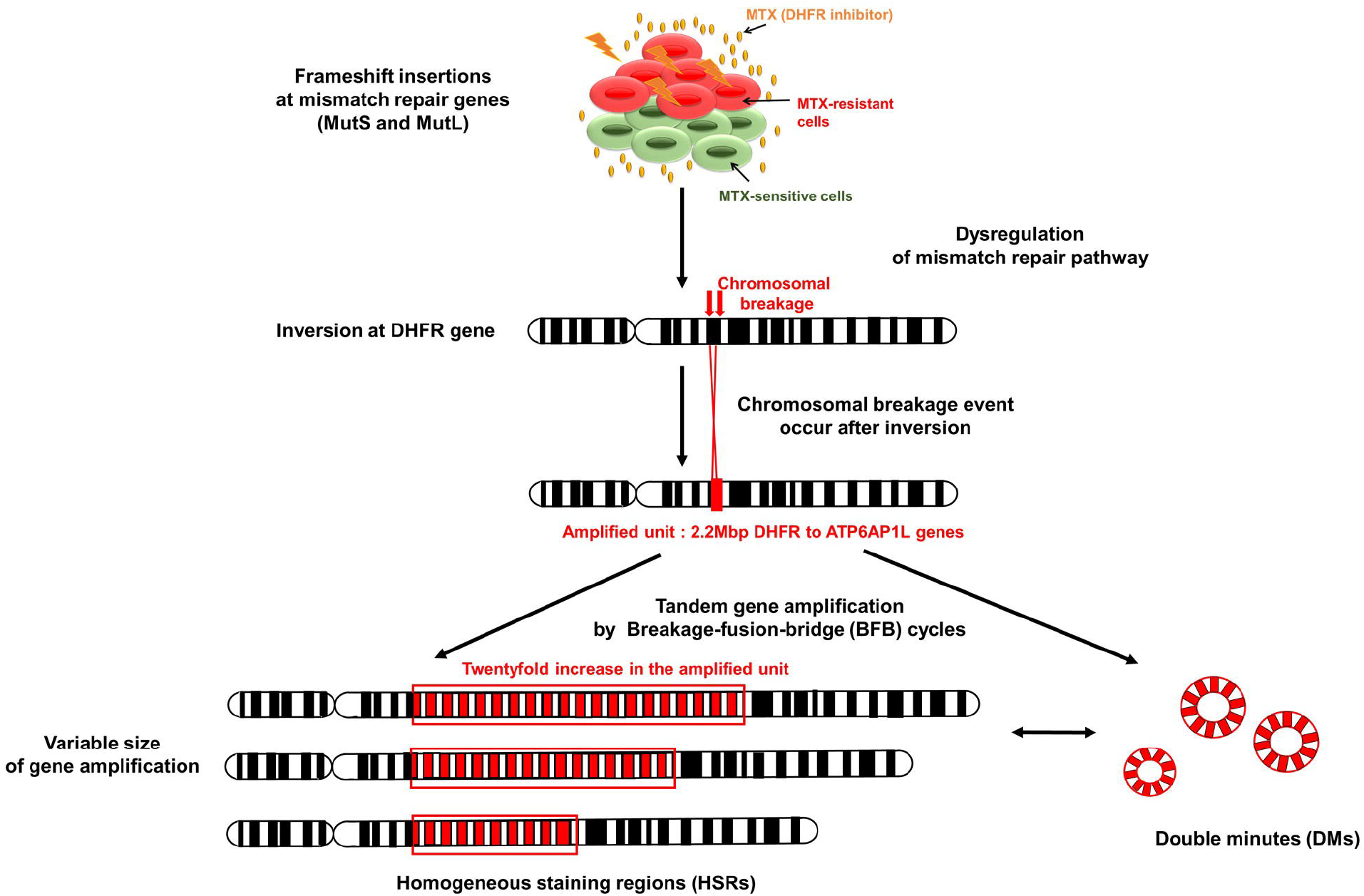
The mechanism of tandem gene amplification in the presence of MTX. Frameshift insertions at the family of mismatch repair genes (mutS and mutL homologs) can initiate the dysregulation of mismatch repair pathways. A chromosomal breakage event occurs after the emergence of an inversion at *DHFR*. Those that occurred between *DHFR* and *ATP6AP1L* (2.2 Mb) were involved in producing the amplified unit. Finally, a variable size of tandem amplifications can be achieved via breakage fusion-bridge (BFB) cycles.

After malfunctioning in mismatch repair mechanisms in the presence of MTX, chromosomal breakages occurred at the *DHFR* gene on chromosome 5, because of the inverted repeat at the start position of the gene. Additionally, inverted repeats from the *DHFR* gene to *ATP6AP1L* (2.2 Mb) were involved in producing the amplified unit. The endpoint of the amplified unit had the same inverted repeat, which could indicate the stop position for gene amplification. However, it is still unknown how the specific genes were selected and involved in gene amplification mechanisms.

Finally, variable sizes of gene amplification with HSRs could be produced via BFB cycles, and the unstable HSRs could be occasionally transformed into either a different size of HSR fragment or DMs, accompanying inversions at the endpoints. Overall, the co-amplification of *MSH3* and *DHFR* as well as frameshift mutations in *MSH* and *MLH* continuously affected genetic instability and enhanced the resistance to methotrexate.

## Discussion

Cancer cells grown in the presence of anti-cancer drugs occasionally have many copies of specific genes, but the molecular mechanisms behind such adaptations are as yet unknown, since analyzing repetitive DNA sequences is limited by many technical difficulties and in evaluating alignment of NGS reads and data processing^46,47^. Recent developments in NGS technologies have enabled large reads (~10Kb) and are able to generate enormous amounts of genomic information at base-level resolution. Such detailed data can facilitate analysis of unknown regions, which is not possible using other technologies^48, 49^.

Here, complicated cancer genomics and abnormal structures were detected and analyzed using a combination of advanced technologies including the PacBio SMRT, optical mapping, and Hi-C analysis, as well as short-read sequencing and FISH. An integrative framework was also used to detect and decipher structural variations in MTX-resistant HT-29 cells. This made use of a large repeat and complex DNA segments. Finally, the amplified region was characterized and evaluated by size and relevant genetic defects, and potential gene amplification mechanisms were suggested.

In order to analyze complicated repetitive sequences involved in gene amplification in the presence of MTX, several MTX-resistant clones were selected and generated from a HT-29 cell line by increasing MTX concentration in a MTX sensitization study. When an amplified *DHFR* gene from several clones was detected via FISH, the cells in each clone had a very large variation of the 5q12 region. *DHFR* was amplified in a small portion of cells, given that tumor cells responded heterogeneously to MTX. Regardless, the effects of the drug would be dissimilar, even if the cells were grown in the same conditions^50, 51^. This could lead to bioinformatics difficulties in finding accurate amplified patterns and repetitive sequences in previous as well as future studies.

In order to overcome the limited analysis provided by heterogeneously amplified patterns*, DHFR* amplified patterns in specific clones were optimized via serial dilution and repeated single-cell selection to generate homogenous amplified patterns. Finally, an optimized clone was obtained that had 96% *DHFR*-amplified cells. Approximately 75% of the cells in the clone converged to a homogeneous *DHFR* amplification pattern.

After optimizing a resistant clone, the amplified unit and tandem gene amplification of 11 genes, from the *DHFR* gene to the *ATP6AP1L* gene on chr5 (2.2Mbp), were identified through the long-range genomic information from long-read sequencing and optical genome mapping. The amplified unit had high coverage (~197X) compared to control (~10X). This implied that the amplified region was approximately twenty-fold tandemly amplified compared to the original sequence, which was confirmed from the high gene expression pattern and alternatively splicing patterns. Gene expression on the amplified unit was highly overexpressed, going from a 5-fold change to a 122-fold change compared to that of the control. These variable expression patterns could be affected by the complexity of splicing patterns, even if these genes were amplified simultaneously.

Furthermore, an inversion at both the start and endpoints of the amplified unit was detected in long-read sequencing data, and this structure variant was cross-validated via optical genome mapping. Previous studies have also shown that an inversion in gene amplification could lead to the initiation of gene amplification by stimulating chromosomal breakage. In addition, such a phenomenon could represent a transformation from circular DNA segment DMs to intra-chromosomal HSRs^52, 53^.

Using Hi-C analysis, we identified high intra-chromosomal interactions on the amplified region and significant interactions at the novel TADs. Long-range chromosomal interactions, from 109Mbp to 138Mb more than those of the amplified region, were also unexpectedly detected. The amplified unit in the Hi-C analysis exactly matched that of the long-range genomic information and RNA sequencing results. This could explain why the amplified position was compactly packed and had frequent contact with each other, which could represent a chromatin architecture for gene amplification^54^.

We also found that the copy number in chromosome 5 was significantly higher in MTX-resistant HT-29 than that of control. Moreover, DMs, in the form of extrachromosomal DNA and another type of chromosomal abnormality, were detected in the amplified region and not in FISH analyses. This could indicate that this structure exists on highly compacted chromosome positions and harbors the amplification of several genes by generating a new chromosomal structure DM and dynamically converting it to another type of chromosomal HSR, which was previously described in neuroblastoma study^55^. This could explain the involvement of chromosomal abnormalities in MTX resistance. However, the mechanism involved in the conversion of HSRs between DMs could not be explained because of the unknown time point of the transition between two structures, its low coverage (10X) in sequencing data, and a large variation in amplification size.

Taken together, these findings suggest that *DHFR* could not be the only target for a MTX-associated mechanism, since 11 unsuspected genes were tandemly amplified between *DHFR* and *ATP6AP1L*, and the expanded region on the 5q arm over the amplified region interacted with other amplified genes each other. The highest gene expression in the amplified unit was *RASGRF2*, which was a 122-fold increase over control. Conversely, *DHFR* underwent a five-fold increase over control. Therefore, more investigations are needed to target and assess the role of *RASGRF2* in MTX related gene amplification mechanism, which might utilize the promoter for *DHFR* to obtain its high efficiency in the presence of MTX.

In addition, novel frameshift insertions in *MSH* and *MLH* were identified in the MTX-resistant sample, which could play an important role in the rapid progression of gene amplification and MTX resistance. In this vein, the microsatellite instability (MSI), i.e., the effect of deficient DNA mismatch repair in colon cancer, was tested to evaluate the status of MMR in MTX-resistant HT-29 cells. There was no significant difference between MTX-resistant cells and control^56^.

This implied that MSI could not affect genetic instability and the entire MMR system over whole chromosomes; whereas, MTX toxicity could induce mutations on MMR genes as well as genetic predisposition on chromosome 5 in MTX-resistant HT-29 cells by inserting adenine or thymine nucleotides on *MSH* and *MLH* genes, thereby enabling the harboring of gene amplification. This points to a possible tandem gene amplification mechanism that progresses through breakage-fusion bridge cycles.

Further studies are needed to identify whether a frameshift insertion in *MSH3* might be caused by co-amplification of *DHFR* and other frameshift insertions that result in the malfunctioning of mutS homologs and hyper-mutability in the presence of significant MTX^57, 58^. However, frameshift insertions in *MSH* and *MLH* could not be generated by the gene amplification process, since all mutated genes except for *MSH3* are located outside of the amplified unit. Therefore, the frameshift mutations in each gene should have had a different cause in the presence MTX toxicity as well as different impacts on MTX-resistant HT-29 cells.

In addition, mismatch repair location is recognized by two mutS homologs such as mutS-alpha (*MSH2* and *MSH6*), which is known for repair of single-nucleotide mismatches, and muS-beta *(MSH2* and *MSH3*), which is known as a repair mechanism for large indels^59, 60^. Such mutS homologs should need mutL homologs (*MLH1* and *PMS2*) for binding to a recognition site.

Therefore, mutations in mutS and mutL homologs could prevent repair from a single-nucleotide mismatch and large indels in MTX-resistant HT-29 cells. The prevention of mutations and inversions in mutS and mutL homologs could increase cells’ sensitivity to MTX and inhibit gene amplification. In the future, methods of preventing these mutations and structural variants could be developed to overcome drug-resistance and guide optimal clinical use of such drugs.

Most of all, additional validation for the identified SVs and repetitive sequences is required to interpret the entire sequence of the gene amplification mechanism and its impact on MTX resistance. This information will provide a clue to how cancers adapt to drugs. In this vein, inversions at the start and end positions on the amplified unit affect chromosomal breakage and formation of DMs from the repetitive sequence. Additionally, the effects of MTX-associated mechanisms on frameshift insertions in *MMR*, *MLH*, and inversions along the amplified region should be described and validated through a comprehensive analysis of the involved genes and pathways, Although several limitations exist in this study, our findings may give new insight into the mechanisms underlying the amplification process and evolution of MTX-resistance in colon cancer and leukemia. Moreover, the use of Hi-C data to detect unforeseen chromosomal rearrangements such as DMs and HSRs has been shown to be promising for future analyses.

## Conclusions

Complex NGS technologies can be used to identify complicated genomic sequences and novel structural variants that are difficult to detect with other technologies. Whenever possible, a comprehensive suite of tools and techniques is required to interpret repetitive sequences and structural variations in gene amplification. This will help to identify the most important therapeutic mechanisms and new targets of anti-cancer drugs, which affect various intracellular pathways across many time and length scales.

Finally, our findings will bolster clinical cancer studies and inform diagnoses as well as the management and treatment of various cancers, and provide in-depth guidance toward pharmacologic targets for anti-cancer drugs as well as personalized medicine.

## Methods

### Preparation and sensitization studies for MTX-resistant cancer cells

After curating a list of cancer cell lines based on previous studies, we targeted and maintained human colon adenocarcinoma HT-29 cell lines from the Korean Cell Line Bank (KLCL) to develop MTX-resistant cancer cells, as described previously^61^. HT-29 was chosen because it can be engineered to grow in high concentrations of MTX via the amplification of *DHFR*.

To generate MTX-resistant HT-29 cells, we optimized the MTX concentration and graded concentrations (from 10^−8^ to 10^−6^ M) in five T25 flasks per cell line, using RPMI media supplemented with 10% fetal bovine serum. The MTX solution involved mixing 100mg MTX powder with 1.967ml DMSO. This mixture was then aliquoted at 100 µl/ml and kept at −20℃. A limited number of resistant cells (~3×10^5^) from each T25 flask was plated in Petri dishes in the same culture medium and MTX concentration.

We then used 3.2mm-diameter cloning discs (Sigma) for 3 isolate clones (C1-2,C4-3,C8-22) from each Petri dish, which were grown up in another T25 flask under the same conditions. Subsequently, MTX-resistant clones were passaged 40 times, without MTX. For the second and third treatment cycles, 3×10^5^ clones and parental HT-29 cells were added and cultured in 25 cm^2^ flasks. A stepwise treatment of MTX concentration from 10^−8^ mol/L to 10^−6^ mol/L was applied; MTX-resistant cells were grown in the 10^−6^ mol/L MTX concentration.

### Detection of gene amplification

The copy number of MTX-resistant clones was measured via TaqMan Copy Number Reference Assays and Viia7 technology to select clones with highly amplified *DHFR* genes. The copy number of the DHFR gene was estimated via relative quantitation (RQ), using the comparative Ct method, and computed using the Ct difference (delta Ct) between MTX-resistant samples and reference samples (Hapmap NA19982). Then, the ∆Ct values of MTX-resistant samples were compared to a Hapmap reference sample known to have two copies of *DHFR*, such that they were two times the relative quantity of the reference. The following three equations were used to compute the copy number from the Ct value.

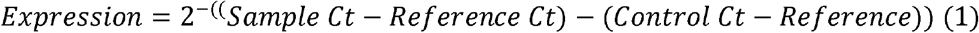

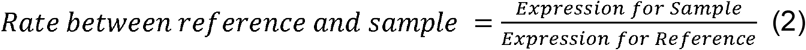

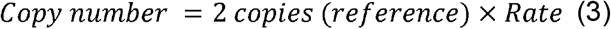

The samples with a high *DHFR* copy number were selected for fluorescence in situ hybridization (FISH) and karyotyping to visualize and map *DHFR*, and detect chromosomal abnormalities. The DHFR red signal and 5p12-green signal were counted at anaphase and metaphase, with 1000x magnification and a triple (RGB) filter.

### RNA-seq analysis and transcriptome profiling

RNA-seq for MTX-resistant HT-29 cells and control samples was performed to investigate expression profiling. Total RNA, with ribosomal RNA removed, was prepared and sequenced using Illumina HiSeq (depth 100X), as previously described^62^. The obtained reads were mapped to the human reference genome (GRCh38) using the Spliced Transcripts Alignment to a Reference (STAR) tool to produce analysis-ready BAM files. We followed the key principles of processing and analysis steps from the GATK website. The mapped reads (BAM file) were visualized via SeqMonk.

To estimate the expression of each gene, raw reads were counted using the HTSeq-count tool and normalized to variance stabilizing data (VSD) expression via the R package DEseq2. Fragments per kilobase million (FPKM) values were calculated using the R package edgeR, and converted to log_2_ values. The median-centered gene expression was computed from FPKM expression, using Cluster 3.0 software, which subtracts the row-wise median from the expression values in each row. The median-centered VSD and FPKM expression were visualized in a heatmap.

### Variant discovery analysis

Variant calling was performed on transcriptome datasets. The duplicated sites from analysis-ready BAM files were filtered with Picard, and variants were called and filtered by removing spurious and known RNA-editing sites, in VCF format. The genomic variants were compared between control and MTX-resistant HT-29 sample. We performed the variant discovery analyses by observing the step-by-step recommendations provided by the Genome Analysis Toolkit (GATK) to obtain high-quality variants^63^.

In order to determine the exact SNPs from the call set, the variants were filtered out according to several conditions. First, the cutoff for quality-by-depth (QD) was 3.0, which is the variant confidence score divided by the unfiltered depth of coverage. Variants were filtered out if they had a score less than 3.0. Second, the variants were filtered out when the Fisher Strand (FS) score was >30.0, which indicated the Phred-scaled p–value using Fisher’s Exact Test for detecting strand bias^64^. The identified and filtered variants were annotated using RefSeq genes and the ANNOVAR tool.

### Differentially expressed gene analysis

Differentially expressed genes between MTX-resistant HT-29 cells and control samples were analyzed using the R packages DESeq2 and edgeR (*P*-value□<□0.05, |Log2 (fold change)| ≥ 1, and baseMean ≥ 100). The differentially expressed genes were determined for enrichment analysis with KEGG gene sets via Gene Set Enrichment Analysis (GSEA).

### Alternative splicing event analysis

The exon inclusion levels, defined with junction reads from RNA sequencing results, were subsequently processed via rMATS.3.2.5^65^. Five different types of alternative splicing event (SE:Skipped exon, MXE: Mutually exclusive exon, A5SS: Alternative 5’ splice site, A3SS: Alternative 3’ splice site, RI: Retained intron) were identified in both MTX-resistant HT-29 cells and control samples. The number of significant events was detected using both junction counts and on-target reads.

### PacBio long-read sequencing analysis

Genomic DNA was extracted from MTX-resistant HT-29 cells and control samples using the Gentra Puregene Cell kit (Qiagen) and the library for PacBio. The PacBio long-reads were aligned to the human genome (version GRCh38) with BWA-mem aligner, and the pre-processing pipeline on the BWA-mem website was followed to ensure technical and biological quality results. The read depth was estimated via the Depth-of-Coverage option in GenomeAnalysisTK. Coverage differences between MTX-resistant HT-29 cells and control bigger than 10 were selected for determining the amplified region.

### Detection of genomic variants and amplification units

The structural variants (deletion, duplication, inverted duplication, translocation, and inversion) in MTX-resistant HT-29 cells and control samples were analyzed using Sniffles from sorted PacBio output derived from BWA-MEM^66^. The BAM files were converted to binned copy numbers across a genome using Copycat. The genomic rearrangements were visualized from VCF files (Sniffles) and read coverage files (Copycat) via SplitThreader (http://splitthreader.com/)^67^.

### Long-read sequencing and optical genome mapping

Optical mapping of the PacBio assembly data was performed using the BioNano Assembler software (Irys System, BioNano Genomics) to get accurate sequences. High molecular weight DNA was isolated using the IrysPrep Plug Lysis Long DNA Isolation Protocol (Bionano Genomics). In brief, cells were trypsinized, washed in FBS/PBS, counted, rewashed in PBS, and embedded in agarose plugs using components from the Bio-Rad Plug Lysis Kit. The plugs were subjected to Proteinase K digestion (2 × 2 h at RT). After a series of washes in buffer from the Bio-Rad kit, followed by washes in TE (Tris-EDTA), the plugs were melted and treated with GELase enzyme (Epicenter).

The high molecular weight DNA was released and subjected to drop dialysis. The DNA was left to equilibrate for 4 d, then quantified using the Qubit Broad Range dsDNA Assay Kit (Thermo Fisher Scientific). Using the IrysPrep NLRS assay (Bionano Genomics), 200–300 ng/µL of high molecular weight DNA underwent single-strand nicking with 10 units of Nt.BspQI nickase (New England BioLabs). Nicked sites were repaired with fluorophore-labeled nucleotide to restore strand integrity.

The backbone of fluorescently labeled double-stranded DNA was stained with the intercalation dye YOYO-1. Labeled molecules were directly loaded onto IrysChip, without further fragmentation or amplification, and imaged using the Irys instrument. Multiple cycles were performed to reach an average raw genome depth-of-coverage of 50x. Additionally, the tandem repeats in the amplified region in both the assembly and raw data were identified using IrysView 2.0 software.

### Hi-C data analysis

Approximately 50 million MTX-resistant HT-29 cells and control cells were used for high-throughput chromatin conformation capture (Hi-C) data sets. We generated 2 Hi-C libraries using the HindIII restriction enzyme, following a previously established protocol^68^. In brief, the Hi-C protocol involves crosslinking cells with formaldehyde, permeabilizing them with nuclei intact, digesting DNA with a suitable restriction enzyme, filling the 5’-overhangs while incorporating a biotinylated nucleotide, ligating the resulting blunt-end fragments, shearing the DNA, capturing the biotinylated ligation junctions with streptavidin beads, and analyzing the resulting fragments with paired-end sequencing via Hi-Seq2000.

The HiC data (fastq files) was processed to normalized contact matrices using HiC-Pro version 2.10.0^69^. The pipeline was based on the bowtie 2 aligner and the selected restriction enzyme (HindIII) was used to generate normalized contact maps, as described in the HiCPro pipline (https://github.com/nservant/HiC-Pro). Each aligned read was assessed to determine the valid interactions and control quality by excluding invalid ligation products and duplicated valid pairs. The aligned Hi-C sam file were converted into the HiCnv format, which calls CNVs from Hi-C data^70^. Also, inter-chromosomal translocations and their boundaries were detected using HiCtrans from a Hi-C matrix file. The list of valid interaction output files called by HiC-pro was converted to a Juicebox input file and visualized using Juicebox (https://github.com/theaidenlab/juicebox/wiki).

The R package HiCcompare was used to detect differential spatial chromatic interactions, on a genome-wide scale, between control and MTX-resistant HT-29 cells^71^. Using this package, the adjusted interaction frequencies after adjustment of joint-normalization functions with the adjusted p-values and filtered low-average expression were represented after multiple testing correction.

### Statistical test

All statistical analyses were performed using R-3.3.0. The gene expression between MTX-resistant HT-29 sample and control (HT-29) was compared, and the p-value was indicated using the unpaired Student’s t-test or Mann-Whitney U-test, based on the Shapiro-Wilk normality test. P-values less than 0.05 were considered to be statistically significant.

## Abbreviations

A3SS: Alternative 3’ splice site
A5SS: Alternative 5’ splice site
BAM: Binary alignment map
BFB: Breakage-fusion-bridge
CNV: Copy number variation
DEG: Differentially expressed gene
DEL: Deletion
DHFR: Dihyrofolate reductase
DUP: Duplication
FDR: False discovery rate
FISH: Fluorescent In Situ Hybridization
FPKM: Fragment per kilo base per million
GATK: Genome analysis toolkit
GSEA: Gene set enrichment analysis
Hi-C: High throughput chromosome conformation capture
IGV: Integrative genomics viewer
INV: Inversion
INVDUP: Inverted duplication
KEGG: Kyoto encyclopedia of genes and genomes
Lsign: Sum of the sign of the entries in the lower triangle
Lvar: the variance of the lower triangle
MTX: Methotrexate
MXE: Mutually exclusive exon
ncRNA: non-coding RNA
NGS: Next generation sequencing
PacBio: Pacific Bioscience
RI: Retained intron
RNA-seq: RNA sequencing
SE: Skipped exon
SMRT: Single molecule real-time
SNV: Single nucleotide variation
Score: Conner score
STAR: Spliced transcripts alignment to a reference
SV: Structural variation
TAD: Topologically associating domains
Usign: Sum of the sign of the entries in the upper triangle
Uvar: The variance of the upper triangle
VSD: Variance stabilizing data
WGS: Whole-genome sequencing

## Data availability

PacBio long read sequencing and whole transcriptome sequencing data are available under the NCBI Sequence Read Archive (SRA) accessions no. SRPXXXXX

## Acknowledgements

This work has been supported by Macrogen Inc. (grant no. MGR17-04)

## Contributions

J.-S.S. and J.-Y.S. conceived and designed the experiments. A.K. conducted the experiment by preparing the MTX resistant HT-29 sample and extracting DNA and RNA. J.-Y.S. performed PacBio long read Seq and whole transcriptome Seq experiments. A.K. performed sequencing data processing, bioinformatics and statistical analyses. J.-S.S., A.K. and J.-Y.S wrote the manuscript

