## Supplementary Figures and Tables for "Genomic and transcriptomic analyses reveal a tandem amplification unit of 11 genes and mutations of mismatch repair genes in methotrexate-resistant HT-29 cells"

### Supplementary Figures 1-14
### Supplementary Tables 1-7

#### Supplementary Figures List

**Supplementary Figure 1. The morphological change of MTX resistant colon cancer cells.**
**Supplementary Figure 2. The detection of the amplified *DHFR* gene at 5q arm in MTX resistant C1-2.**
**Supplementary Figure 3. The visualization of the amplified *DHFR* gene at 5q arm in MTX resistant C1-2.**
**Supplementary Figure 4. The detected FISH patterns of amplified *DHFR* gene in MTX resistant C1-2.**
**Supplementary Figure 5. The visualization of inter-chromosomal genomic rearrangements in MTX resistant HT-29 and control samples.**
**Supplementary Figure 6. The multiple-read view of alignment over amplified region with Ribbon.**
**Supplementary Figure 7. The scaffolding of PacBio long reads over amplified region compared to hg 38.**
**Supplementary Figure 8. Genome mapping over the amplified region.**
**Supplementary Figure 9. The expression level in 5q 14.2 region.**
**Supplementary Figure 10. The difference of non-synonymous mutations between MTX resistant HT-29 and control over whole chromosomes.**
**Supplementary Figure 11. Genome-wide view of intra-chromosomal interactions.**
**Supplementary Figure 12. The topologically associating domains (TADs) on chromosome 5.**
**Supplementary Figure 13. The comparison of relative copy number between MTX resistant and control samples.**
**Supplementary Figure 14. Schematic workflow.**

#### Supplementary Tables List

**Supplementary Table 1. The estimation of copy number and expression of *DHFR* gene in MTX resistant clone (C1-2) at each cycle.**
**Supplementary Table 2. The detected variants on chromosome 5 in MTX resistant HT-29.**
**Supplementary Table 3. The comparison of mutations between control and MTX resistant HT-29 over whole chromosomes.**
**Supplementary Table 4. The comparison of mutations between control and MTX resistant HT-29 on chromosome 5.**
**Supplementary Table 5. The detection of novel frameshift insertions in MTX resistant HT-29 compared to control.**
**Supplementary Table 6. The differentially expressed genes (DEGs) in MTX resistant HT-29.**
**Supplementary Table 7. The topologically associating domains (TADs) with high intra-chromosomal interactions on chromosome 5.**

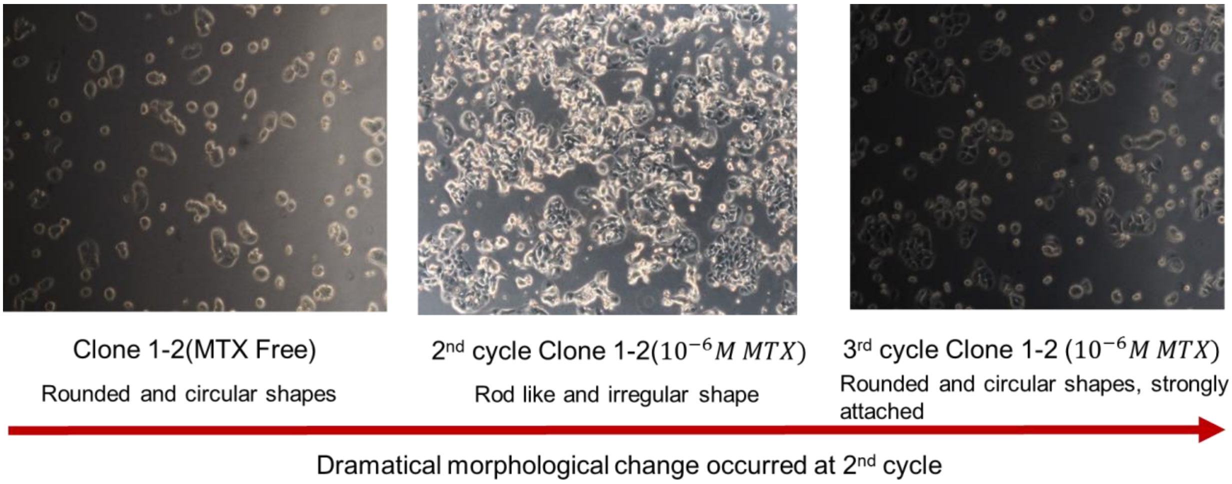


**Supplementary Figure 1. The morphological change of MTX resistant colon cancer cells.** The MTX resistant clone (C1-2) was detected and captured by light microscope at 200x magnification. The morphological changes at each cycle were indicated under $10^{-6}$ mol/L MTX.

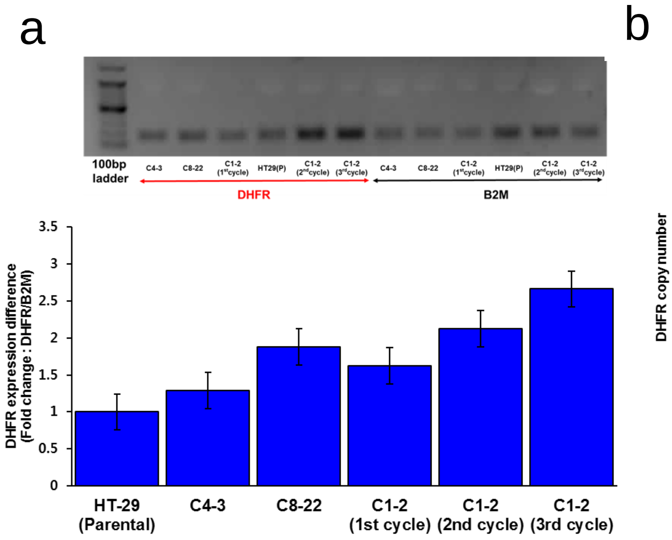


**Supplementary Figure 2. The detection of the amplified *DHFR* gene at 5q arm in MTX resistant C1-2.** The expression (a) and copy number (b) of *DHFR* gene in MTX resistant clones and C1-2 clone at each cycle were estimated by qPCR. The error bars indicated standard error (SE) of the mean.

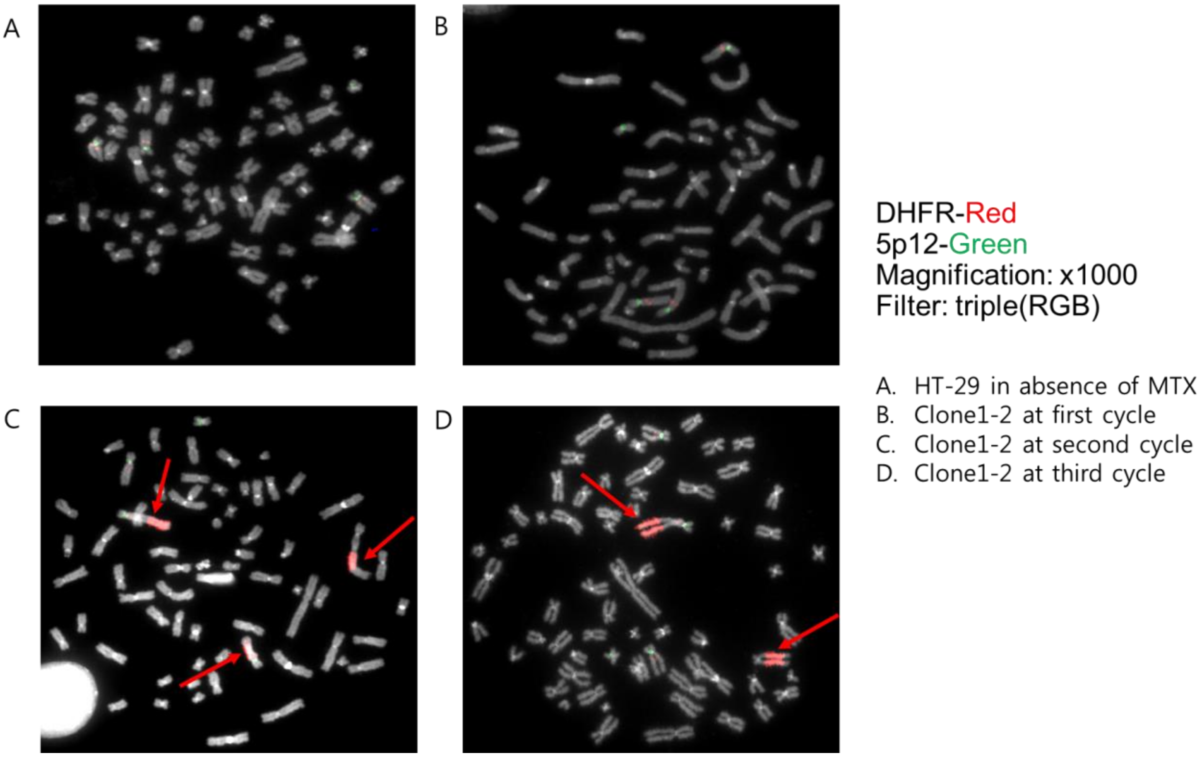


**Supplementary Figure 3. The visualization of the amplified *DHFR* gene at 5q arm in MTX resistant C1-2.** The amplified DHFR gene was visualized by Fluoscent In Situ Hybridization (FISH) on MTX resistant clone (C1-2) at each cycle and control sample with the 1000x magnification. The amplified *DHFR* gene region was indicated by the red arrow.

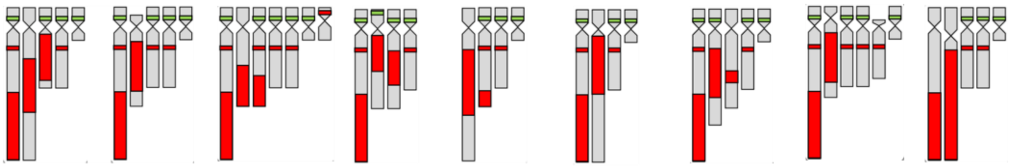

| FISH Signal Type | Type 1 | Type 2 | Type 3 | Type 4 | Type 5 | Type 6 | Type 7 | Type 8 | Type 9 |
|---|---|---|---|---|---|---|---|---|---|
| DHFR-Red Signal count | P3,S2 | P2,S3 | P3,S5 | P3,S2 | P2,S2 | P2,S2 | P3,S2 | P2,S4 | P2,S2 |
| 5p12-Green signal count | 4 | 4 | 5 | 4 | 3 | 3 | 4 | 4 | 3 |
| Metaphase count (%) | 11 (44) | 7 (28) | 1 (4) | 1 (4) | 1 (4) | 1 (4) | 1 (4) | 1 (4) | 1 (4) |

**Supplementary Figure 4. The detected FISH patterns of amplified *DHFR* gene in MTX resistant C1-2.** The FISH signal type of amplified DHFR gene in MTX resistant clone (C1-2) was displayed with the paining signal (P) and spot signal (S). 5q12 was indicated by green signal, and *DHFR* gene was indicated by red signal. Each number of red and green signal and metaphase were counted.

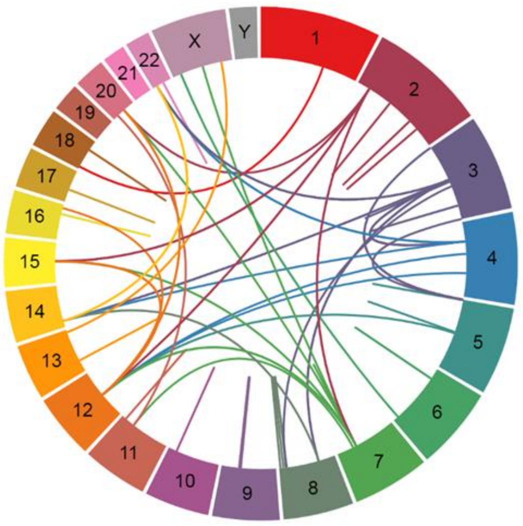


Control (HT29)

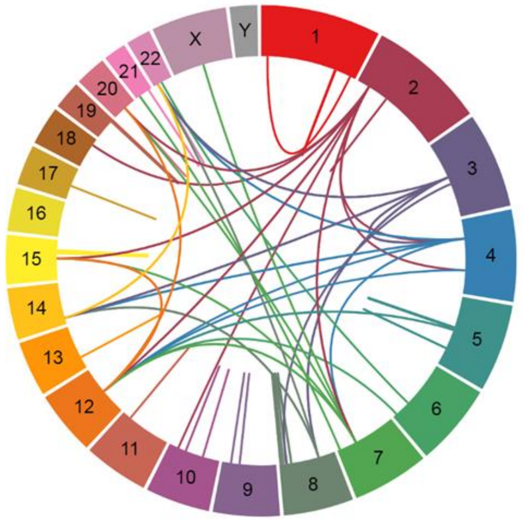


MTX resistant HT29

**Supplementary Figure 5. The visualization of inter-chromosomal genomic rearrangements in MTX resistant HT-29 and control samples.** The inter-chromosomal genomic rearrangements were compared between MTX resistant HT-29 and control samples, and it was visualized by Splitthreader over all chromosomes.

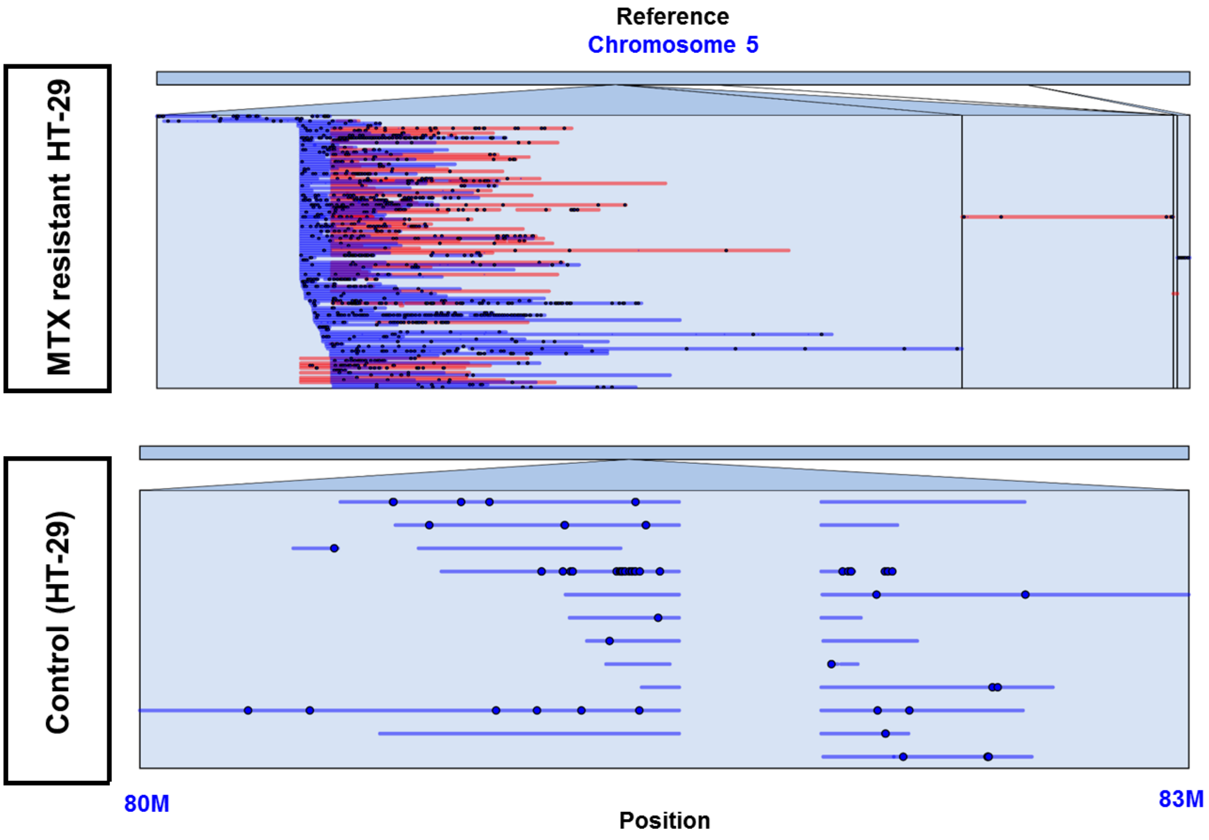


**Supplementary Figure 6. The multiple-read view of alignment over amplified region with Ribbon.** The multi-reads from alignment data (BAM) was visualized and compared between control and MTX resistant sample on chromosome 5: 80,000,000-83,000,000 by using Ribbon. Blue and red lines indicated the forward and reverse strands, respectively.

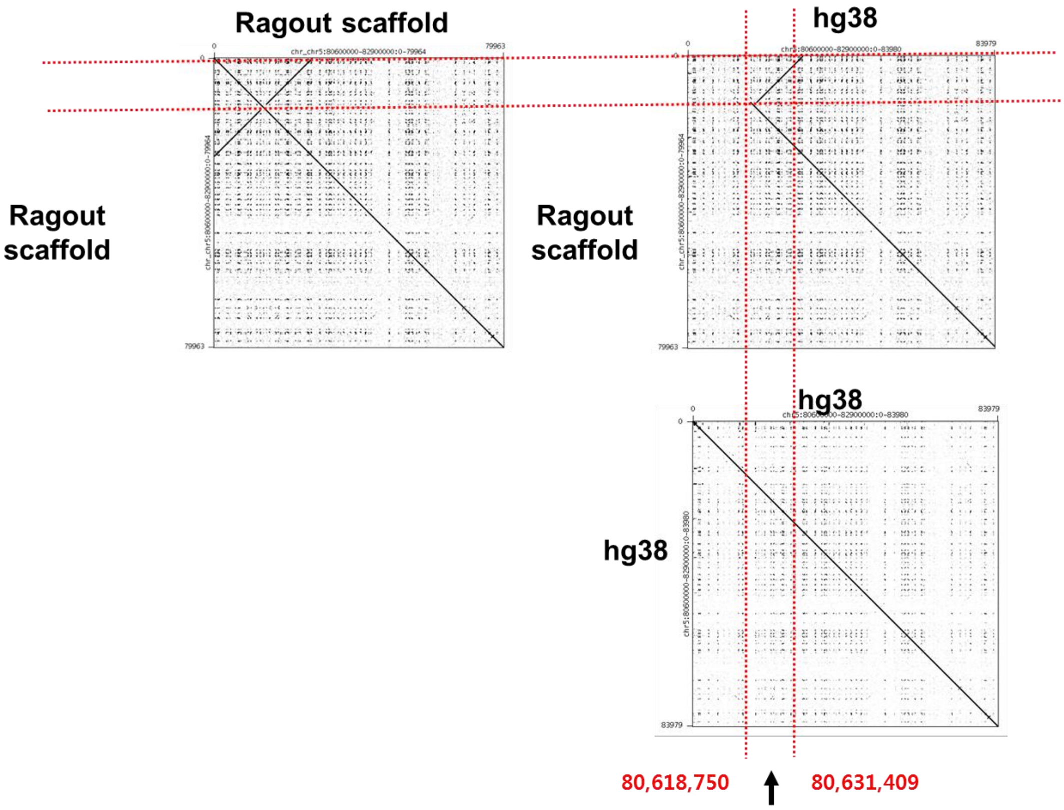


**Genes in inverted region**

| | |
|---|---|
| LINC01337 | long intergenic non-protein coding RNA 1337 |
| DHFR | Homo sapiens dihydrofolate reductase |
| CTC-325J23.2 | antisense RNA |
| MTRNR2L2 | Homo sapiens MT-RNR2-like 2 |

**Supplementary Figure 7. The scaffolding of PacBio long reads over amplified region compared to hg 38.** The inversion at the start point of duplication unit was visualized by the three dot plots compared to hg38.The inverted region and its genes were indicated with the description.

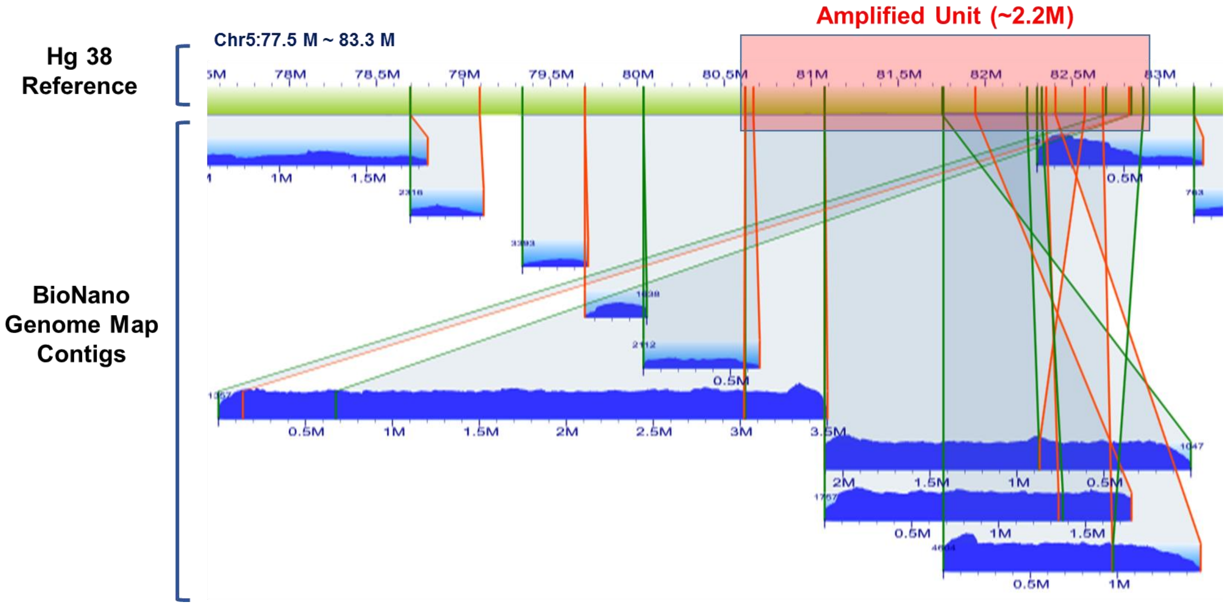


**Supplementary Figure 8. Genome mapping over the amplified region.** The BioNano genome map contigs around amplified region were visualized and compared along with the reference (hg38). The start position of contig indicated with the green color, and the end position of contig indicated with the orange color.

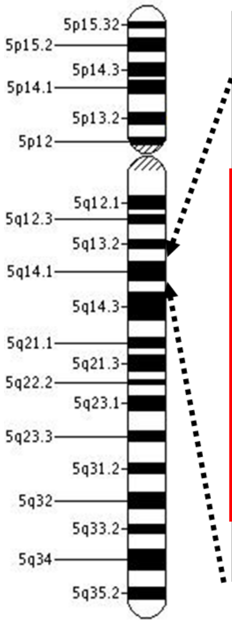


| Gene | Expression (FPKM) | | | | |
|---|---|---|---|---|---|
| | Control | MTX-resistant sample | Foldchange | Log2foldchange | Difference |
| ZFYVE16 | 4.83 | 4.89 | 1.01 | 0.02 | 0.06 |
| FAM151B | 0.11 | 0.17 | 1.55 | 0.63 | 0.06 |
| ANKRD34B | 0 | 0 | 1.00 | 0.00 | 0.00 |
| DHFR | 32.04 | 174.13 | 5.43 | 2.44 | 142.09 |
| MTRNR2L2 | 1.69 | 10.18 | 6.02 | 2.59 | 8.49 |
| MSH3 | 6.46 | 139.41 | 21.58 | 4.43 | 132.95 |
| RASGRF2 | 4.72 | 578.83 | 122.63 | 6.94 | 574.11 |
| CKMT2 | 0.7 | 41.1 | 58.71 | 5.88 | 40.40 |
| ZCCHC9 | 4.48 | 202.69 | 45.24 | 5.50 | 198.21 |
| ACOT12 | 0 | 0.84 | - | - | 0.84 |
| SSBP2 | 0.6 | 4.15 | 6.92 | 2.79 | 3.55 |
| ATG10 | 99.14 | 1382.08 | 13.94 | 3.80 | 1282.94 |
| RPS23 | 149.25 | 2028.95 | 13.59 | 3.76 | 1879.70 |
| ATP6AP1L | 0.31 | 14.19 | 45.77 | 5.52 | 13.88 |
| TMEM167A | 34.86 | 10.19 | 0.29 | -1.77 | -24.67 |
| XRCC4 | 7.12 | 0 | 0.17 | -2.59 | -5.94 |

**Supplementary Figure 9. The expression level in 5q 14.2 region.** The table depicted the expression level (FPKM) over duplicated region, and the fold change and difference of expression were computed between control and MTX resistant HT-29. The fold change, which is bigger than 5, was indicated with the red box. .

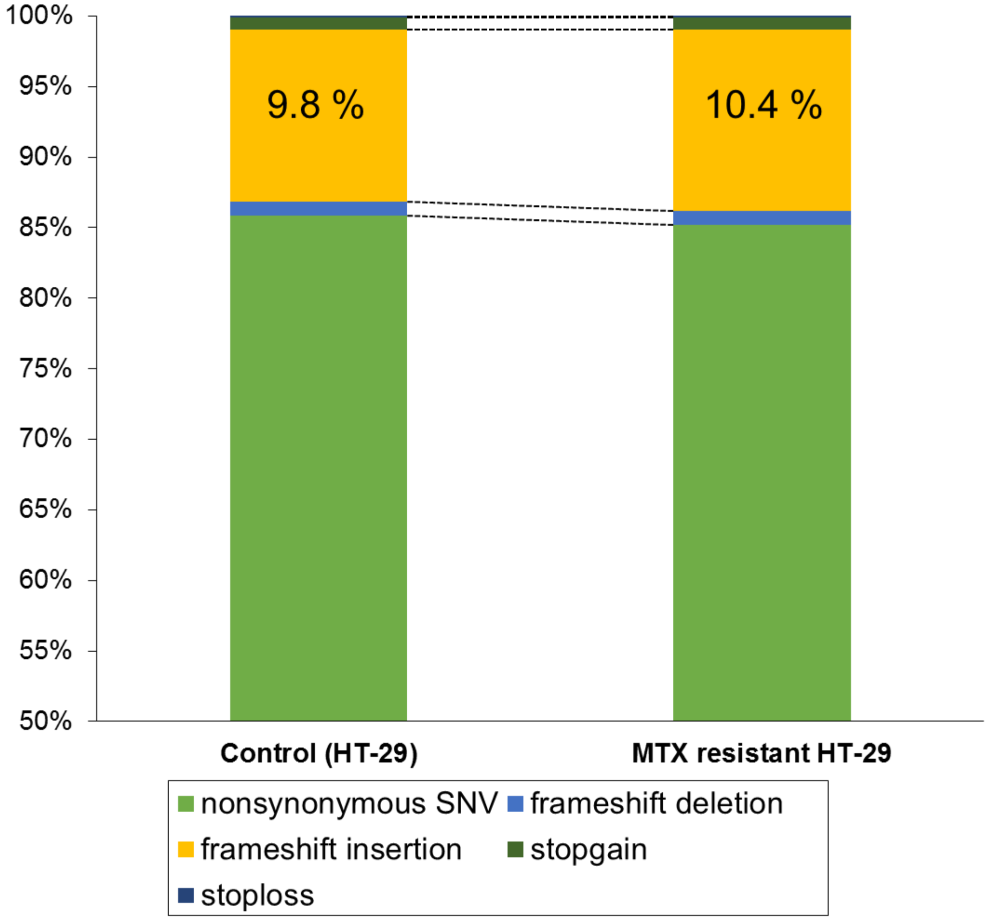


**Supplementary Figure 10. The difference of non-synonymous mutations between MTX resistant HT-29 and control over whole chromosomes.** The non-synonymous mutations (non-synonymous SNV, frameshift deletion, frameshift insertion, stop-gain, and stop-loss) were compared between MTX resistant HT-29 and control over whole chromosomes, and the percentage of frameshift insertions was indicated.

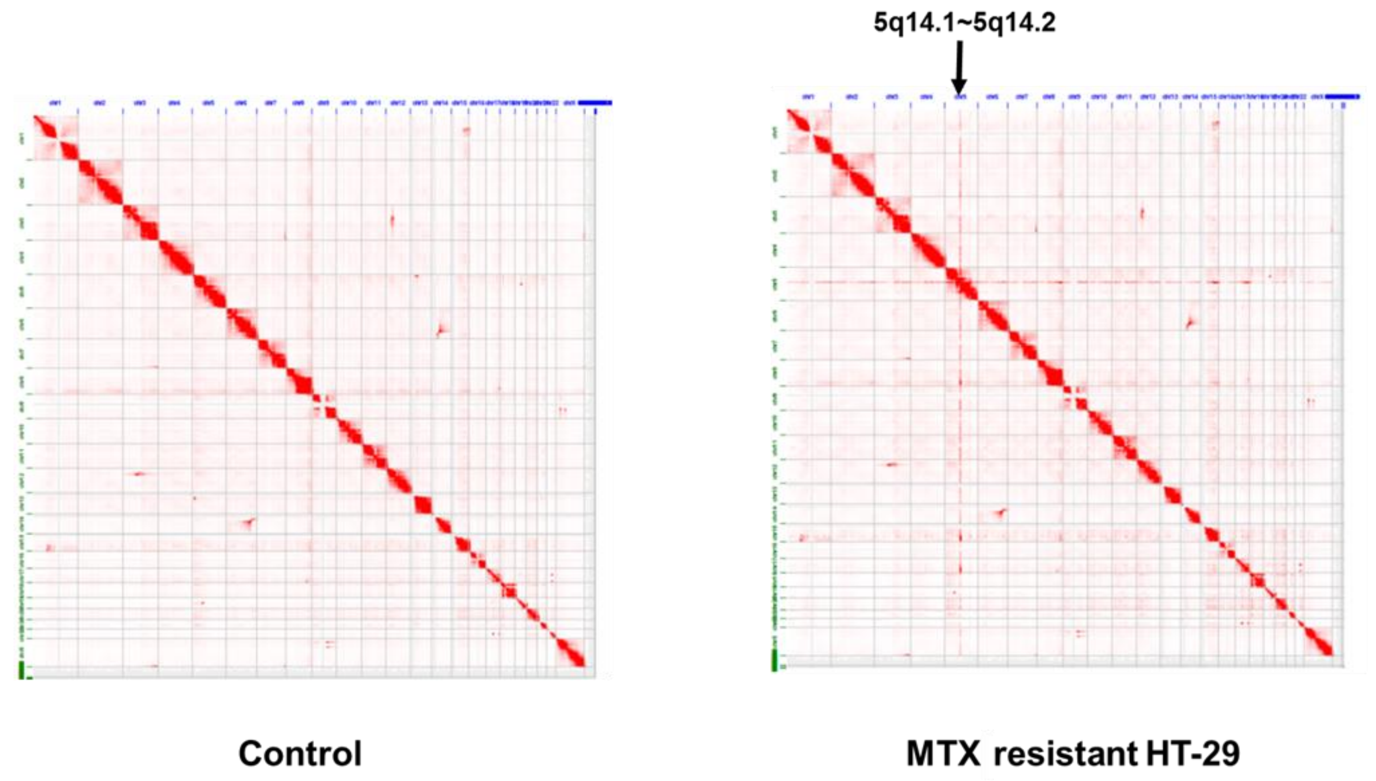


**Supplementary Figure 11. Genome-wide view of intra-chromosomal interactions.** The intra-chromosomal interactions over genome-wide view in MTX resistant HT-29 and control were visualized by Juicebox at 5kb resolution, and the amplified region was indicated by the arrow.

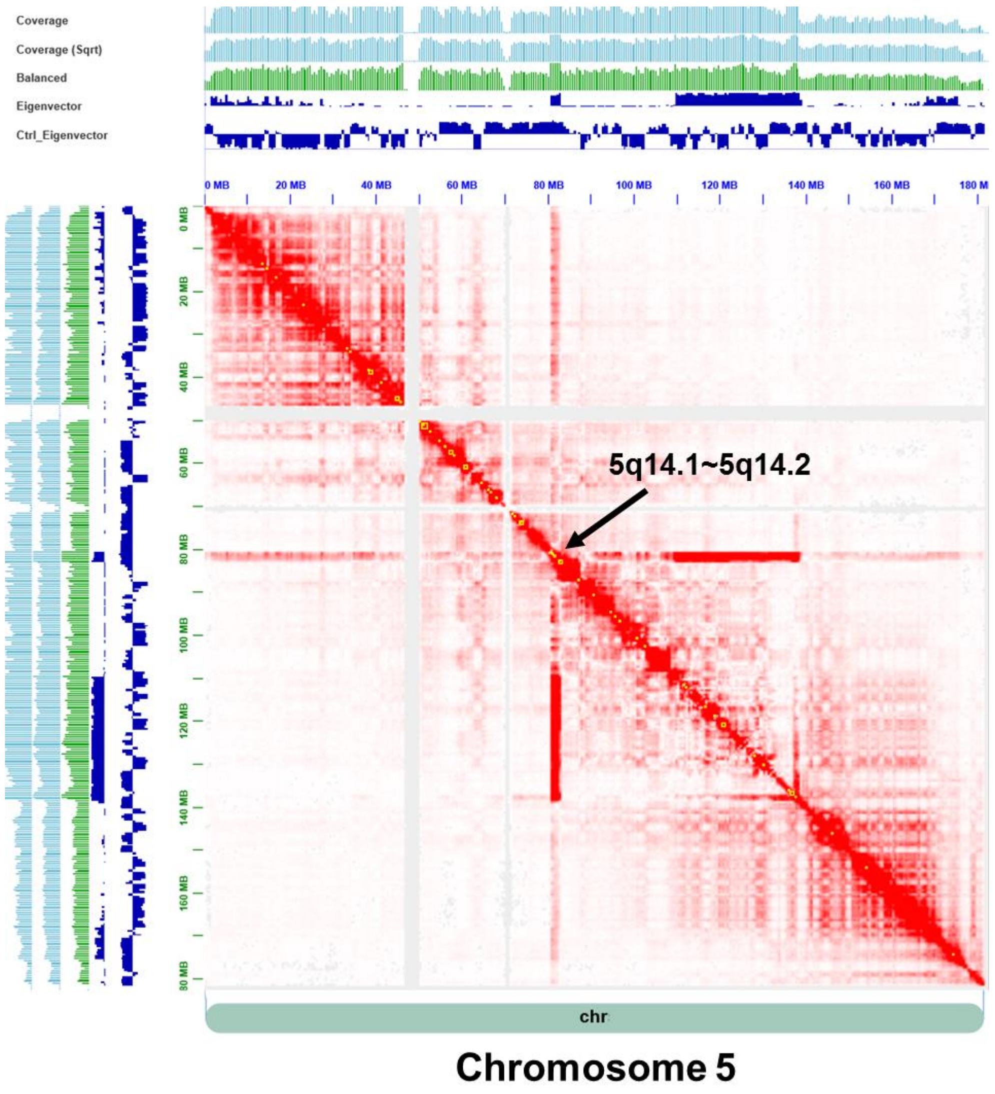


**Supplementary Figure 12. The topologically associating domains (TADs) on chromosome 5.** The topologically associating domains were identified by Arrowhead algorisms and visualized with the intra-chromosomal interactions on chromosome 5. The TADs were yellow-boxed.

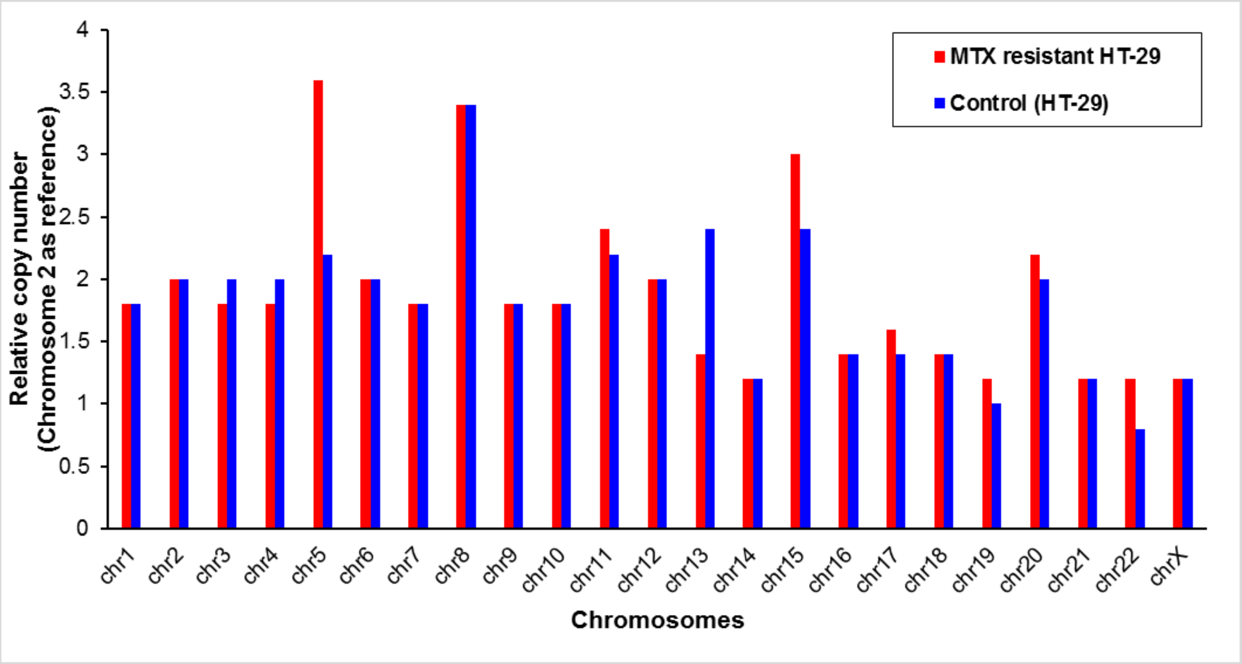


**Supplementary Figure 13. The comparison of relative copy number between MTX resistant and control samples.** The relative copy number over whole chromosomes was estimated by HiCnv from Hi-C data. The copy number of chromosome 2 was used as the reference.

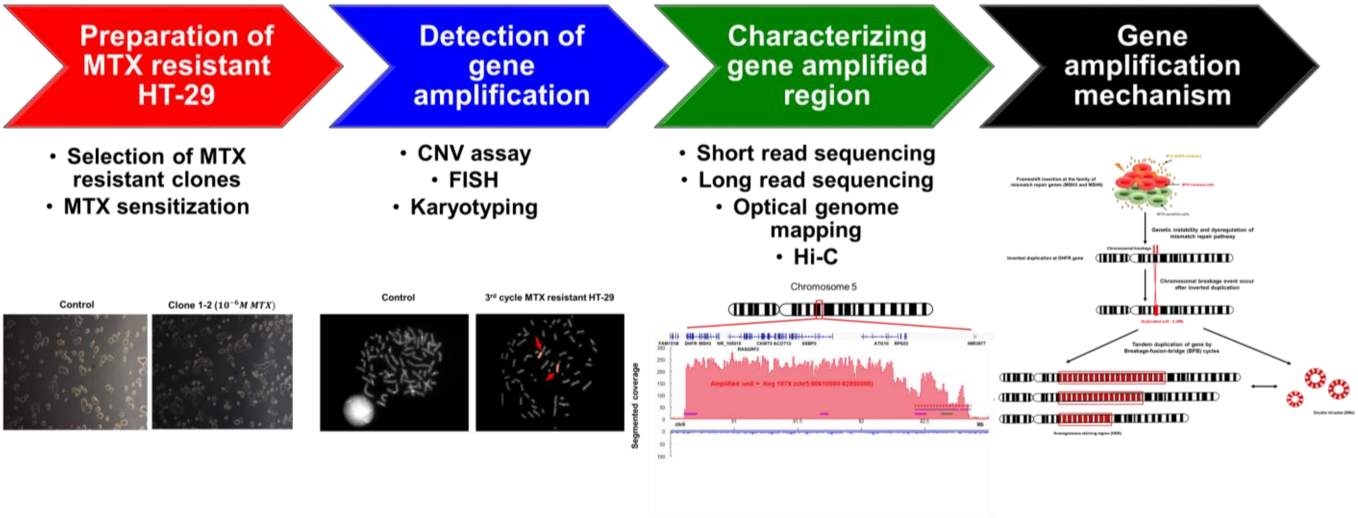


**Supplementary Figure 14. Schematic workflow.** The experimental and computational analyses were performed on methotrexate resistant colon cancer cell line (HT-29) in order to characterize the gene amplified region and understand the underlying mechanism.

**Supplementary Table 1. The estimation of copy number and expression of *DHFR* gene in MTX resistant clone (C1-2) at each cycle.**

| Type of Sample | Sample Ct | Expression | Rate for Copy Number | Copy Number |
|---|---|---|---|---|
| Hapmap NA19982 | 24.99 | 2.52 | 1.00 | 2.00 |
| Control (HT-29) | 26.33 | 1.00 | 0.40 | 0.79 |
| 1st cycle MTX resistant HT-29 | 26.04 | 1.22 | 0.48 | 0.97 |
| 2nd cycle MTX resistant HT-29 | 20.21 | 69.17 | 27.42 | 54.83 |
| 3rd cycle MTX resistant HT-29 | 19.99 | 80.56 | 31.93 | 63.87 |

#### Supplementary Table 2. The detected variants on chromosome 5 in MTX resistant HT-29

| chrom1 | pos1 | strand1 | chrom2 | pos2 | strand2 | variant_name | variant_type | split | size | CNV_category | category | nearby_variant_count |
|---|---|---|---|---|---|---|---|---|---|---|---|---|
| 5 | 80617089 | - | 5 | 80618750 | - | 24 | INVDUP | 47 | 1661 | matching | simple | 0 |
| 5 | 81781592 | + | 5 | 81781648 | + | 50 | INVDUP | 10 | 56 | neutral | simple | 0 |
| 5 | 82245207 | + | 5 | 82573904 | + | 56 | INV | 56 | 328697 | matching | crowded | 1 |
| 5 | 82470789 | - | 5 | 82842464 | + | 58 | DUP | 11 | 371675 | matching | crowded | 4 |
| 5 | 82470887 | + | 5 | 82804501 | + | 59 | INV | 12 | 333614 | partial | crowded | 4 |
| 5 | 82473031 | - | 5 | 82474560 | - | 60 | INVDUP | 18 | 1529 | matching | simple | 3 |
| 5 | 82678370 | + | 5 | 82764579 | - | 62 | DEL | 56 | 86209 | matching | simple | 3 |
| 5 | 82804502 | + | 5 | 82842464 | + | 64 | INV | 47 | 37962 | matching | simple | 3 |
| 5 | 137519108 | + | 5 | 137520622 | - | 65 | DEL | 19 | 1514 | neutral | simple | 0 |
| 5 | 137686889 | + | 5 | 137688197 | - | 66 | DEL | 22 | 1308 | neutral | simple | 0 |
| 5 | 88112313 | - | 12 | 66057592 | - | 256 | TRA | 12 | -1 | neutral | reciprocal | 12 |
| 5 | 88112381 | + | 12 | 66057683 | + | 284 | TRA | 12 | -1 | neutral | reciprocal | 12 |

**Supplementary Table 3. The comparison of mutations between control and MTX resistant HT-29 over whole chromosomes.**

| Mutation types | MTX resistant HT-29 | Control (HT-29) |
|---|---|---|
| Exonic total | 13982 | 13310 |
| Exonic splicing | 10 | 9 |
| Splicing | 70 | 155 |
| ncRNA exonic | 6040 | 5876 |
| ncRNA exonic:splicing | 2 | 3 |
| ncRNA splicing | 21 | 26 |
| 3'UTR | 28523 | 27892 |
| 5'UTR | 3448 | 3632 |
| Nonsynonymous SNV | 5708 | 5730 |
| Frameshift deletion | 70 | 66 |
| Frameshift insertion | 858 | 816 |
| Stopgain | 57 | 55 |
| Stoploss | 9 | 9 |

**Supplementary Table 4. The comparison of mutations between control and MTX resistant HT-29 on chromosome 5.**

| Mutation types | MTX resistant HT-29 | Control (HT-29) |
|---|---|---|
| Exonic total | 577 | 559 |
| Exonic splicing | 1 | 0 |
| Splicing | 4 | 8 |
| ncRNA exonic | 159 | 206 |
| ncRNA exonic:splicing | 0 | 0 |
| ncRNA splicing | 1 | 1 |
| 3'UTR | 1332 | 1337 |
| 5'UTR | 166 | 164 |
| Nonsynonymous SNV | 220 | 226 |
| Frameshift deletion | 4 | 3 |
| Frameshift insertion | 51 | 12 |
| Stopgain | 6 | 4 |
| Stoploss | 0 | 0 |

### Supplementary Table 5. The detection of novel frameshift insertions in MTX resistant HT-29 compared to control.

| MTX resistant HT-29 | | | | | | | | | |
|---|---|---|---|---|---|---|---|---|---|
| **Chr** | **Start** | **End** | **Ref** | **Alt** | **Func.refGene** | **Gene.refGene** | **Gene Title** | **ExonicFunc.refGene** | **ExAC_ALL** |
| chr2 | 47803552 | 47803552 | - | T | exonic | MSH6 | mutS homolog 6 | frameshift insertion | . |
| chr2 | 189818083 | 189818083 | - | A | exonic | PMS1 | postmeiotic segregation increased 1 | frameshift insertion | 3.54E-05 |
| chr2 | 189863838 | 189863838 | - | A | exonic | PMS1 | postmeiotic segregation increased 1 | frameshift insertion | . |
| chr3 | 37012077 | 37012077 | A | G | exonic | MLH1 | mutL homolog 1 | nonsynonymous SNV | 0.2325 |
| chr5 | 80654962 | 80654962 | A | G | exonic | MSH3 | mutS homolog 3 | nonsynonymous SNV | 0.9029 |
| chr5 | 80675095 | 80675095 | - | A | exonic | MSH3 | mutS homolog 3 | frameshift insertion | . |
| chr5 | 80854162 | 80854162 | A | G | exonic | MSH3 | mutS homolog 3 | nonsynonymous SNV | 0.8731 |
| chr5 | 80873118 | 80873118 | G | A | exonic | MSH3 | mutS homolog 3 | nonsynonymous SNV | 0.7305 |
| chr7 | 5987144 | 5987144 | T | C | exonic | PMS2 | postmeiotic segregation increased 2 | nonsynonymous SNV | 0.8514 |
| chr7 | 5987357 | 5987357 | G | A | exonic | PMS2 | postmeiotic segregation increased 2 | nonsynonymous SNV | 0.3854 |
| chr7 | 5987525 | 5987525 | - | T | exonic | PMS2 | postmeiotic segregation increased 2 | frameshift insertion | . |
| chr14 | 75047125 | 75047125 | G | A | exonic | MLH3 | mutL homolog 3 | nonsynonymous SNV | 0.4126 |
| chr14 | 75047180 | 75047180 | T | C | exonic | MLH3 | mutL homolog 3 | nonsynonymous SNV | 0.9968 |
| **Control (HT-29)** | | | | | | | | | |
| **Chr** | **Start** | **End** | **Ref** | **Alt** | **Func.refGene** | **Gene.refGene** | **Gene Title** | **ExonicFunc.refGene** | **ExAC_ALL** |
| chr3 | 37012077 | 37012077 | A | G | exonic | MLH1 | mutL homolog 1 | nonsynonymous SNV | 0.2325 |
| chr5 | 80654905 | 80654905 | - | CCGCAGCGC | exonic | MSH3 | mutS homolog 3 | nonframeshift insertion | 0.0427 |
| chr5 | 80654962 | 80654962 | A | G | exonic | MSH3 | mutS homolog 3 | nonsynonymous SNV | 0.9029 |
| chr5 | 80854162 | 80854162 | A | G | exonic | MSH3 | mutS homolog 3 | nonsynonymous SNV | 0.8731 |
| chr5 | 80873118 | 80873118 | G | A | exonic | MSH3 | mutS homolog 3 | nonsynonymous SNV | 0.7305 |
| chr7 | 5987144 | 5987144 | T | C | exonic | PMS2 | postmeiotic segregation increased 2 | nonsynonymous SNV | 0.8514 |
| chr7 | 5987357 | 5987357 | G | A | exonic | PMS2 | postmeiotic segregation increased 2 | nonsynonymous SNV | 0.3854 |
| chr14 | 75047125 | 75047125 | G | A | exonic | MLH3 | mutL homolog 3 | nonsynonymous SNV | 0.4126 |
| chr14 | 75047180 | 75047180 | T | C | exonic | MLH3 | mutL homolog 3 | nonsynonymous SNV | 0.9968 |

**Supplementary Table 6. The differentially expressed genes (DEGs) in MTX resistant HT-29.**

| | |
|---|---|
| Up-regulated DEGs (383 genes) | PRSS22 BAIAP3 ETV7 CD22 ALOX5 TYMP BTN3A1 USP2 TRAF1 SLC2A3 CACNB1 GPR116 CLEC2D TRIB2 SPEG SIDT1 GSDMB DHRS9 SCARF1 CACNG4 RGS11 P2RX5 COL16A1 HSD17B14 ASAP3 EPB41L1 TMEM40 SLC4A11 LAG3 ARHGAP4 ANKRD24 YPEL3 PITPNM2 MYO15A NLRP1 DFNB31 CATSPERG CYP2D6 APOL4 UPK3A REC8 WFDC2 MAP1LC3A ATP11C KCND1 VGLL1 GABRE GDPD3 BMF RHOV MAP4K1 AMH IL4I1 DENND3 CCDC114 PBX4 SH3GL2 UNC5B MAP3K8 RASD1 DHX58 MYH3 MGP ADTRP ULBP1 MSH3 RASGRF2 HES1 SLC4A3 PCSK4 FN1 MLPH PADI2 GBP1 SGK1 CTGF TRPM6 LTBP2 GPR68 DUSP1 CD274 PLEKHG1 TP53AIP1 EGR1 TNFSF10 EGR2 NR4A1 C4BPB ARHGAP40 PMEPA1 ZBP1 PRICKLE4 CDKN1A RUNX2 C20orf195 IL1B C3 FOSB ID1 PRKCG ZFP36 APOL3 APOL2 APOBEC3F KRT17 LOXL1 CHRNA10 APOE TNNI2 PNPLA7 ANGPTL6 C19orf66 PLXNA3 SH3BP5 PDLIM4 CKMT2 ZCCHC9 LGALS3 TRIM22 XAF1 REEP2 ALDH3B2 FBXO44 EPSTI1 RARRES3 RSAD2 SLC37A2 ANXA1 ELF5 EGR4 WNT10A THSD1 SCEL RTP4 CASP1 IFI44L MYOF LOXL4 DUSP5 GPR87 ADAMTS14 SEMA7A RASGEF1B ITGB7 ITGAX ZMYND15 ATHL1 SIK1 CACNG8 PADI1 SYTL1 CYR61 CTSK REN CSRNP1 KLHL3 SERPING1 LYPD6B IL18 ATG10 ACOXL NR4A2 GPR110 ANKRD29 CERS3 GOLGA7B UBE2L6 TNFRSF14 DUSP2 XDH BTG2 CCDC17 PAQR6 FOXH1 SLC25A34 SCNN1D GBP4 ATF3 IVL ELF3 IFI16 KIAA1407 ZC3H12A SHROOM1 TRPV6 OTUD1 DKFZP686J19100 HDGFRP3 ANKDD1A CERCAM PRRT2 KIFC2 RILP TRPV3 TMEM88 TRANK1 TTC21A MFSD7 THBS3 CXCL10 IL8 TMEM154 FOS CST1 STAT2 CEL KRT13 EFEMP2 SYT12 C11orf80 TDRD12 SLC26A9 SLC22A1 DDIT3 VWA3A CST6 UCP3 IDSP1 SLCO3A1 C8orf31 SAMD9L JUN ODF3B ALS2CL EGR3 HCLS1 MUC16 HEPHL1 BAI1 UBA7 LINC00085 HCAR2 MX2 EMILIN3 KCNQ3 TACSTD2 FAM70B DUSP8 SOCS3 IFNE MAFF HEATR7B1 SOCS1 MAPK11 STAC3 MUC1 C11orf35 TLCD2 PDIA2 CYP4F12 RPS23 CEACAM19 MIR22HG EPOR MAGI2 DNAJB13 PALM3 HES4 NOXA1 APOD C6orf222 CXCL17 CGB7 HSH2D C5AR1 BCO2 CCDC154 SLC22A20 ZNF44 PLCG2 ARC GPRASP1 SNORA31 DDO AGER C6orf25 AC096670.3 IGFL4 GPR20 FAM221B LRRC10B FAM71F2 SAP25 ATP6AP1L CPT1B KRTAP5-1 LCAT LBH UCA1 PLIN5 RP11-83B20.1 RP11-429J17.8 RP11-1036E20.9 EVPLL SMTNL1 GCGR hsa-mir-6080 ATAD3C RP11-465B22.3 AC110619.2 SNORD63 ADM5 SH3BP5-AS1 RP11-288L9.4 NTF4 AC009950.2 RP11-203J24.8 RP11-69I8.3 RAB11FIP1P1 SAPCD1 RP4-583P15.10 AC019349.5 GOLGA6L5 RPS23P8 RP11-34A14.3 RP11-263K19.4 RP11-67C2.2 CTD-2020K17.3 RP11-250B2.3 SAPCD1-AS1 RP11-73M7.6 NRADDP CBR3-AS1 RNF223 RP3-430N8.8 AF011889.2 BX470102.3 APOBEC3G PSMB9 UGT1A9 RP11-147I3.1 RP11-379B18.5 AF011889.5 ARHGDIG Metazoa_SRP PDCL3P4 IFITM10 SCARF2 RP3-330M21.5 CTC-281B15.1 CTD-2248H3.1 RP11-510N19.5 CTC-325J23.2 CTD-2193P3.2 HMGB1P3 ALDH1L1-AS1 CTD-2193P3.1 CTC-459I6.1 RP11-459E5.1 CHKB-CPT1B RP11-496I9.1 RP11-326C3.2 MSH5-SAPCD1 RP11-167N4.2 RP11-512N21.3 RP11-631N16.2 RP11-386M24.3 MC1R RP4-647C14.3 RP11-566K11.4 RP11-521C20.3 RP11-1100L3.8 RP11-254F7.2 RP5-1085F17.3 RP11-386M24.6 AC145291.2 RP11-250B2.5 RP11-448G15.3 SPON1 LA16c-325D7.1 LINC00672 SNORD3A RP1-37N7.1 RP13-104F24.3 AC010761.10 RP5-906A24.2 MYO15B AC004510.3 SH3GL1P3 CTD-2240E14.4 CTD-3252C9.4 AC006262.5 CTD-2521M24.5 |
| Down-regulated DEGs (287 genes) | CFTR SLC7A2 PLXND1 USH1C DPEP1 SLC38A5 GRAMD1B HSD17B6 MIPEP ZIC2 KITLG LTBP1 KCNQ1 GYG2 SOAT1 TMEM48 TBXAS1 SLC12A2 IPO5 PRR11 LIMS2 HMMR MCM2 PPP2R2C ENO1 PLD1 GPC4 EPB41L2 SLC4A4 SMARCA2 ABCB1 ORC1 IPO11 TPX2 TESC BIRC5 LYZ DPYSL2 TMEM38B BAMBI PCSK5 LGALS2 C14orf105 SERPINA4 MYBL2 HNF4A FERMT1 FGF9 SLC25A15 CPPED1 KIAA1199 MCM4 CDK6 CPVL COBL MOGAT3 AGR2 SLC1A1 AMBP DKK1 MAP2K6 ANXA10 GALNTL4 SOX6 POU2AF1 CHPT1 UST PERP PRLR HGD PRKAR2A ECT2 HHLA2 PLSCR4 IGFBP2 IL1R2 LEPR AGMAT NR5A2 PROX1 PLAGL1 AKAP7 MYB SLC16A7 GDA ONECUT2 MOB3B CCDC170 LYPLA1 RPL21 FAM126A TUBA1B EREG FOXA2 TMX4 AMOT WNK4 DLGAP5 SGPP1 TSPAN8 TUBA4A VIL1 GNAI1 MGAT3 HOXD13 DLL4 PPP1R1B NR0B2 DDC RFC3 ZDHHC8P1 HMGCS2 LDHA HNF1A AGT SLC19A3 SERPINE2 SMC2 HMGA1 CASC5 SLC38A4 LGR5 FAM222A SLC7A1 BRCA2 DIAPH3 ZIC5 NKD1 IMPA2 GATA6 PRKACB PARP1 LBR CHAC2 CDCA7 PTPRG CCNA2 FAM105A SYTL5 TACC1 LACTB2 FAM171A1 SLC43A1 ADRA2A C11orf53 SLC7A11 SLC2A13 SACS SPC25 CNKSR3 KCTD15 UBASH3B PLCL2 PPARGC1B SLC26A2 PTDSS1 HKDC1 ARSE ZNF618 EDA KALRN FGFR4 SSTR5 AKR7A3 CAMK2N1 PKDCC ARL5A KCNJ3 IHH FABP1 SAP30 MAD2L1 SPINK1 FABP5 HNF4G CLDN2 CDX2 KBTBD6 CKB ANPEP CCDC103 RRM1 FAM83B FEN1 NPNT CXXC4 SLC38A11 GJB1 ROBO1 P2RY1 MYO7B ALCAM SERPINA6 FOXA3 TMEM37 ESCO2 LGALS4 CYCS ADH6 AGR3 CSPG4 MUC13 FBXO45 KCNE3 KBTBD11 C18orf56 GSG2 LCN15 DNAJC22 KCTD12 TMEM64 OR51E1 C3orf58 C8orf33 EPHB3 NEB ASCL2 BRI3BP SFTPA2 TARSL2 GPRIN3 NCR3LG1 MAOA CTSE RP11-57C13.3 SERPINA1 CYP2B6 SPN DPP4 S100A10 PRIM1 AKR1B10 RYR2 PAPSS2 F5 SUCNR1 GRK5 C2orf72 SP5 C10orf112 ZBTB10 LGR4 LDHAP4 MUC5AC TENM3 AC007405.2 SLC25A5-AS1 NPY6R RP1-37C10.3 RP11-103C16.2 AC007163.6 RP11-229P13.23 COL4A2-AS1 AF196970.3 AC013463.2 MYB-AS1 AC005550.3 MLK7-AS1 RPS2P5 RP11-64D22.2 TDGF1 HNF1A-AS1 RP11-67L3.5 RP11-410D17.2 RP11-382J12.1 ADH1C SLC7A11-AS1 RP11-710F7.2 RP11-115D19.1 RAD21-AS1 PRKDC CTD-2292P10.2 RP11-363E6.3 RP11-700F16.3 RP11-627G23.1 RP11-173P15.3 TMPO-AS1 RP11-186F10.2 RP11-210N13.1 RP11-386G11.10 RP11-109D20.2 LINC00261 CTD-2196E14.5 RP11-401P9.4 CTD-2510F5.4 ZNF488 RP11-627G18.1 |

**Supplementary Table 7. The topologically associating domains (TADs) with high intra-chromosomal interactions on chromosome 5.**

| chr1 | x1 | x2 | chr2 | y1 | y2 | score | uVarScore | lVarScore | upSign | loSign |
|---|---|---|---|---|---|---|---|---|---|---|
| chr5 | 80970000 | 81100000 | chr5 | 80970000 | 81100000 | 0.634 | 0.047 | 0.058 | 0.599 | 0.522 |
| chr5 | 81205000 | 81295000 | chr5 | 81205000 | 81295000 | 0.944 | 0.017 | 0.035 | 0.633 | 0.722 |
| chr5 | 81315000 | 81405000 | chr5 | 81315000 | 81405000 | 0.853 | 0.038 | 0.006 | 0.700 | 0.611 |
| chr5 | 81430000 | 81735000 | chr5 | 81430000 | 81735000 | 0.701 | 0.094 | 0.103 | 0.593 | 0.503 |
| chr5 | 81510000 | 81720000 | chr5 | 81510000 | 81720000 | 0.982 | 0.045 | 0.080 | 0.803 | 0.639 |
| chr5 | 81765000 | 81840000 | chr5 | 81765000 | 81840000 | 0.776 | 0.051 | 0.144 | 0.609 | 0.516 |
| chr5 | 82645000 | 82835000 | chr5 | 82645000 | 82835000 | 1.389 | 0.084 | 0.113 | 0.647 | 0.884 |
| chr5 | 82765000 | 82835000 | chr5 | 82765000 | 82835000 | 0.888 | 0.016 | 0.169 | 0.518 | 0.607 |
| chr5 | 82905000 | 83605000 | chr5 | 82905000 | 83605000 | 0.219 | 0.435 | 0.341 | 0.411 | 0.419 |
| chr5 | 87025000 | 87390000 | chr5 | 87025000 | 87390000 | 1.347 | 0.314 | 0.319 | 0.581 | 0.655 |
| chr5 | 90840000 | 91440000 | chr5 | 90840000 | 91440000 | 0.737 | 0.354 | 0.392 | 0.414 | 0.439 |
| chr5 | 94970000 | 95085000 | chr5 | 94970000 | 95085000 | 1.354 | 0.075 | 0.119 | 0.688 | 0.521 |
| chr5 | 95895000 | 96340000 | chr5 | 95895000 | 96340000 | 1.020 | 0.343 | 0.290 | 0.498 | 0.535 |
| chr5 | 96740000 | 97155000 | chr5 | 96740000 | 97155000 | 1.037 | 0.327 | 0.308 | 0.402 | 0.537 |
| chr5 | 100565000 | 101370000 | chr5 | 100565000 | 101370000 | 0.851 | 0.378 | 0.316 | 0.457 | 0.406 |
| chr5 | 102545000 | 103145000 | chr5 | 102545000 | 103145000 | 1.206 | 0.332 | 0.337 | 0.537 | 0.484 |
| chr5 | 102615000 | 103030000 | chr5 | 102615000 | 103030000 | 0.924 | 0.331 | 0.312 | 0.423 | 0.451 |
| chr5 | 111505000 | 112140000 | chr5 | 111505000 | 112140000 | 1.311 | 0.323 | 0.323 | 0.466 | 0.660 |
| chr5 | 112160000 | 112475000 | chr5 | 112160000 | 112475000 | 1.194 | 0.294 | 0.331 | 0.651 | 0.417 |
| chr5 | 116555000 | 116960000 | chr5 | 116555000 | 116960000 | 0.665 | 0.271 | 0.290 | 0.410 | 0.459 |
| chr5 | 118980000 | 119445000 | chr5 | 118980000 | 119445000 | 0.730 | 0.350 | 0.408 | 0.474 | 0.483 |
| chr5 | 120770000 | 121930000 | chr5 | 120770000 | 121930000 | 0.675 | 0.366 | 0.394 | 0.401 | 0.468 |
| chr5 | 128205000 | 128880000 | chr5 | 128205000 | 128880000 | 0.979 | 0.367 | 0.318 | 0.439 | 0.536 |
| chr5 | 130210000 | 130740000 | chr5 | 130210000 | 130740000 | 0.673 | 0.357 | 0.339 | 0.408 | 0.413 |
| chr5 | 136250000 | 136775000 | chr5 | 136250000 | 136775000 | 0.507 | 0.321 | 0.421 | 0.403 | 0.460 |
| chr5 | 136880000 | 137585000 | chr5 | 136880000 | 137585000 | 0.615 | 0.321 | 0.307 | 0.476 | 0.432 |
| chr5 | 137650000 | 137740000 | chr5 | 137650000 | 137740000 | 1.380 | 0.106 | 0.090 | 0.567 | 0.656 |
| chr5 | 146455000 | 146515000 | chr5 | 146455000 | 146515000 | 0.985 | 0.045 | 0.131 | 0.500 | 0.500 |
| chr5 | 154005000 | 154410000 | chr5 | 154005000 | 154410000 | 0.873 | 0.370 | 0.355 | 0.484 | 0.478 |
| chr5 | 174395000 | 174835000 | chr5 | 174395000 | 174835000 | 0.967 | 0.369 | 0.339 | 0.411 | 0.425 |

*Abbreviations*
Score = corner score
Uvar = the variance of the upper triangle
Lvar = the variance of the lower triangle
Usign = -1*(sum of the sign of the entries in the upper triangle)
Lsign = sum of the sign of the entries in the lower triangle
